# An extended Kalman filter for large-volume path positioning of aquatic animals within acoustic telemetry arrays

**DOI:** 10.64898/2026.09.17.752304

**Authors:** James Adam Campbell, Jelger Ellings, Petter Lundberg, Rachel Mawer, Ine Pauwels, Franz Hölker

## Abstract

Acoustic telemetry is a core methodology for collecting fine-scale movement data for aquatic animals. When telemetry receivers are set up in closely spaced arrays with overlapping detection ranges, the detection times of a tagged animal can be used to estimate its position and movement paths. In practice, estimating these paths can be challenging. Traditional time-difference-of-arrival methods generally provide positioning accuracy too poor for inferring fine-scale behaviours, while more robust state-space positioning models can be computationally intensive and practically infeasible to run on large datasets. Here, a novel telemetry positioning method is presented where time-of-arrival positioning is implemented as a state-space model within an extended Kalman filter. The resulting model, termed EK-TOA, provides closed-form solutions to track estimation. Simulated datasets of fish movement within a 2D telemetry array are used to verify the models performance and a real case study is provided where EK-TOA is utilized for the long-term tracking of a tagged fish. In comparison to currently available positioning models, EK-TOA provides a fast and accurate solution for tracking fine-scale movement behaviours of aquatic animals over long, continuous periods of time.

## 2 Introduction

Over the past decade, the fine-scaled analysis of *in situ* aquatic animal movement has been facilitated by the availability of high-resolution underwater acoustic telemetry equipment (Lennox et al. 2023). Acoustic tags attached to animals can emit frequent, short signals on the order of seconds apart, encoding the unique identity of the individual. When acoustic receivers are arranged close together in an array, the locations of these animals can be estimated from the time-of-arrival of these emissions across the array. The resulting movement path can be further analysed with animal movement statistics, providing inference on how environmental or experimental conditions affect the behaviour states and navigation of the animal (Hooten et al. 2017). However, telemetry path positioning — the estimation of these movement paths — can be challenging. As acoustic signals travel fast underwater, small errors in time-of-arrival measures can result in poor positioning accuracy and precision.

Time-difference-of-arrival (TDOA) methods are standard for telemetry positioning and are commonly employed by commercial positioning services (Lennox et al. 2023); they require only detection times and the locations of the receivers in the array (see Smith and Abel 1987). In practice, however, the detection error is often too large to produce suitable tracks for behavioural inference.

Recently, the use of state-space models for underwater telemetry positioning has been popularized by YAPS (yet another positioning solver, Baktoft et al. 2017). State-space models, such as YAPS, supplement measurement data with assumptions about the animals underlying movement processes, giving position estimates that are orders of magnitude more precise than TDOA positioning (Vergeynst et al. 2020). YAPS treats tag positions and emission times as latent states. For a given latent state, the likelihood is evaluated by both the measured detection times across the array — the measurement component — and the assumption that the animal’s movements follow a Gaussian random walk — the process component. YAPS fits these latent states within a maximum likelihood, mixed-model framework using the template model builder (TMB) R package (Kristensen et al. 2016).

While YAPS can provide robust positioning estimates, its performance may be affected by the challenges associated with fitting a large number of latent states via an iterative optimization routine. First, YAPS requires initial values for all tag positions. While these can be randomly sampled positions, providing good initial values can speed up processing times. In particularity challenging setups, good initial values can be necessary to return position estimates. Determining such values, particularity in sparse telemetry arrays where TDOA positioning gives poor results, can be very difficult. Second, the optimization routine is computationally expensive to run. Positioning tracks with many thousands of tag emissions can become time-consuming as the processing time may not scale linearly with the volume of latent states to fit. Additionally, providing too much data can cause the optimizer to fail due to memory limitations.

For large-volume datasets — many thousands of tag emissions captured over days or weeks — YAPS requires segmenting detections data into small periods with manageable processing times. Furthermore, as the accuracy of a given position estimate is tied to the uncertainty of those positions immediately before and after it, positions near the start and end of these track segments tend to have high error. Thus, track segments should ideally be overlapping, causing a further increase in the processing workload.

This paper presents an alternative state-space model for time-of-arrival positioning which addresses the volume and processing time limitations inherent to iterative maximum likelihood routines, such as those used by YAPS, removing the need to segment large datasets. Rather than using an iterative optimizer, the proposed model is formulated so tag positions and emission times can be fitted with an extended Kalman filter (Gelb et al. 1974). As extended Kalman filters provide closed-form solutions to state estimates in sequence, large-volume tracks can be positioned near-instantaneously and processing times scale linearly with the volume of data. The resulting time-of-arrival model (hereafter referred to as EK-TOA) provides position estimates with a comparable quality to those from YAPS. Following the description of the model, simulated data is used to verify the model’s performance. Lastly, an example large-volume case study is presented along with practical guidance on how to use the model.

## 3 Methods

### 3.1 State-space model

State-space positioning combines a measurement model with a process model to estimate the most likely state (i.e. position) of the animal. Here, the state during emission *i* is represented by the vector

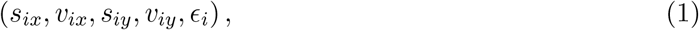

where *s*_*ix*_ and *s*_*iy*_ are the coordinates of the tag emission (i.e. the source), *v*_*ix*_ and *v*_*iy*_ are the instantaneous velocity of the tagged animal, and *ϵ*_*i*_ is an error term used to estimate the tag emission time. Note that for simplicity, we are formulating the state variables in 2-dimensional space. The model can easily be extended to three-dimensional positioning by adding a z-dimension to the relevant following equations.

Given a set of state values, the measurement model returns the time-of-arrival likelihood. The detection time for emission *i* at receiver *k* is given by

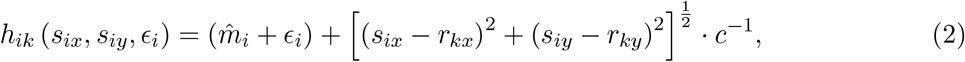

where *r*_*kx*_ and *r*_*ky*_ are the coordinates of the receiver and *c* is the speed of transmission through the water. The true emission time is represented by 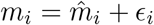, where 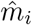 is a prior estimate of the tag emission time and *ϵ*_*i*_ is the error of that estimate.

Often, acoustic telemetry tags are set to emit signals on random uniform intervals, spanning many seconds. Moreover, composite tags are often employed, where multiple signals on unique intervals are emitted in parallel from the same animal. By formulating the true emission time as an error measure of some prior guess, more precise emission time predictions can be provided and no prior knowledge of the tag emission schedule is required. Additionally, the error term *ϵ*_*i*_ can be treated as a Gaussian distribution, allowing fitting within an extended Kalman filter framework.

Extended Kalman filters use the Jacobian of non-linear differentiable functions as linear approximations to those functions. Here, the Jacobian of (2) is given by

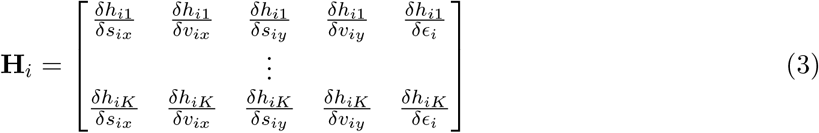

for an array containing *K* receivers with detections where

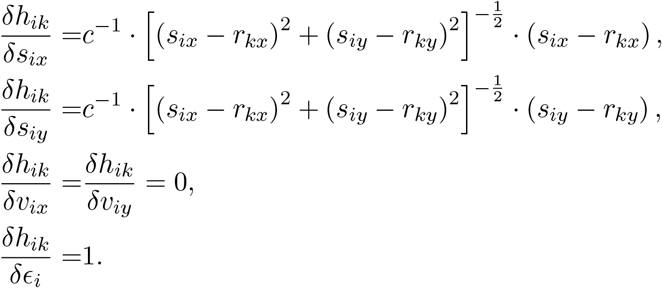

Note that the tag velocity state parameters, *v*_*ix*_ and *v*_*iy*_, are unused by the time-of-arrival model (2).

To model the animals underlying movement — the process model — Johnson et al.’s (2008) continuous-time correlated random walk (CTCRW) model is used. This model treats the animal’s instantaneous velocity along a single dimension as an Ornstein–Uhlenbeck process (see Hooten et al. 2017)

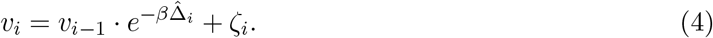

The position process is then given by

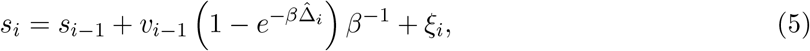

where *ξ*_*i*_ and *ζ*_*i*_ are Gaussian distributed error terms with the corresponding variance/covariance

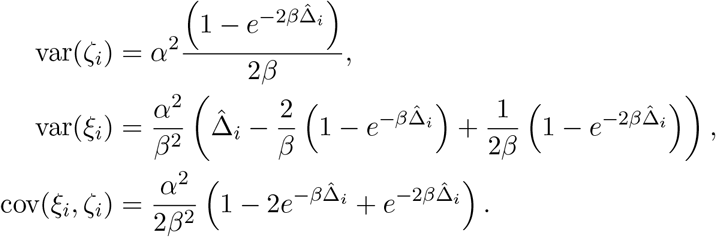

The parameter *β* controls the relaxation rate where lower values assume more ‘momentum’ in the movement of the animal. The time since the last tag emission is given by 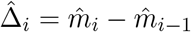.

Correlated walk models are appealing for animal behaviour studies as the smoother tracks can potentially produce more distinguishable behaviour states. As the position and velocity components of the CTCRW model are already linear, they can be arranged in a two-dimensional state transition matrix

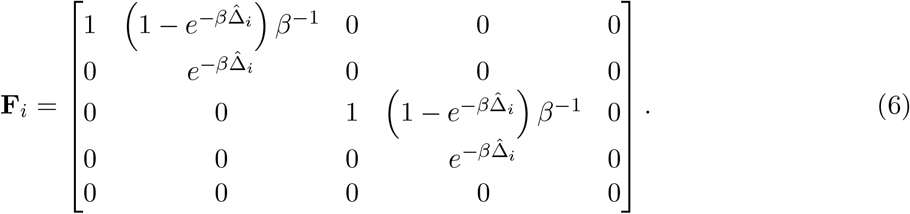

Next, the Gaussian error covariance matrices need to be defined. The observation error covariance is

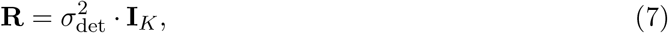

where *σ*_det_ is the standard deviation of the detection error on any receiver and **I**_*K*_ is an identity matrix with dimensions matching the number of receivers with detections in the array. The process error covariance is given by

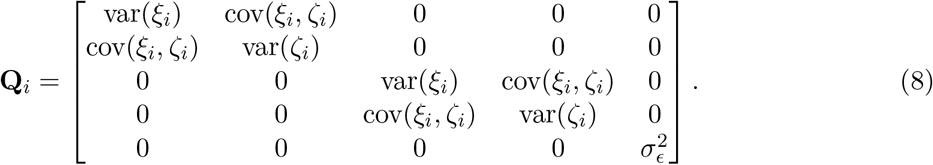

The above equations can be used in an extended Kalman filter. Equations (2) and (3) give the non-linear observation model and its Jacobian, (6) provides the state transition matrix, while (7) and (8) are the observation and process error covariances, respectively. Note that the observation and process parameters *σ*_det_, *σ*_*ϵ*_, *α*, and *β* are constant across all emissions. For those receivers with missing detections, the associated rows and columns in the matrices **H**_*i*_ and **R** can be removed. Receivers with missing data will then be ignored during the update step of the filter for that emission.

Note that the above model can take an alternative formulation where the emission time error, *ϵ*_*i*_, is omitted from the state vector in (1) and instead an emission time covariance matrix is added to the observation error in (7). In the current form, including emission time error within the state vector allows for the simple retrieval of the tag emission time estimates.

Before fitting, some prior estimate of the tag emission times, 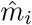, must be provided. This is used in both the time-of-arrival observation model and to roughly estimate the time interval since the previous emission, 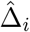. For an agnostic approach, 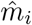 values can be set to each emission’s time of first detection in the array, along with an appropriately large value for *σ*_*ϵ*_. For example, 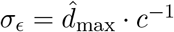, where 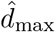 is the farthest distance a tag emission is expected to propagate.

The remaining model parameters (*σ*_det_, *α*, and *β*) can be fitted with maximum likelihood. However, it may be preferable to assign sensible values to these *a priori*. First, many telemetry arrays are equipped with sentinel tags — tags placed at known locations and often with known emission times. Such tags can be used to measure the detection error of the array, providing empirical values for *σ*_det_. Second, the detection error is often too large to facilitate the disentanglement of the CTCRW parameters, *α* and *β*, from the measurement error, *σ*_det_. Lastly, even in the case when detection error is small, an animal can exhibit multiple movement behaviours — expressed as time-varying values for CTCRW parameters. Fitting a single state behaviour model, such as the CTCRW, to multi-state movement data may result in some behaviours being poorly represented in the resulting tracks.

Reasonable *a priori* parameters for the CTCRW model can be chosen by reformulating *α* and *β* into more intuitive measures: the standard deviation of the stationary velocity

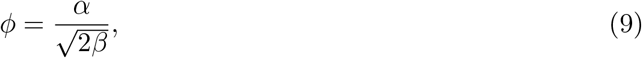

and the time until the velocity correlation falls below 0.05

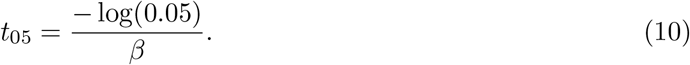

When considering movement in two-dimensional space, the magnitude of the stationary velocity in the horizontal plane follows a Rayleigh distribution with a mode of *ϕ* and a mean of *ϕ*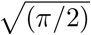. Figure 1 shows the relationship between *ϕ* and the resulting stationary horizontal velocity distribution.

**Figure 1.**
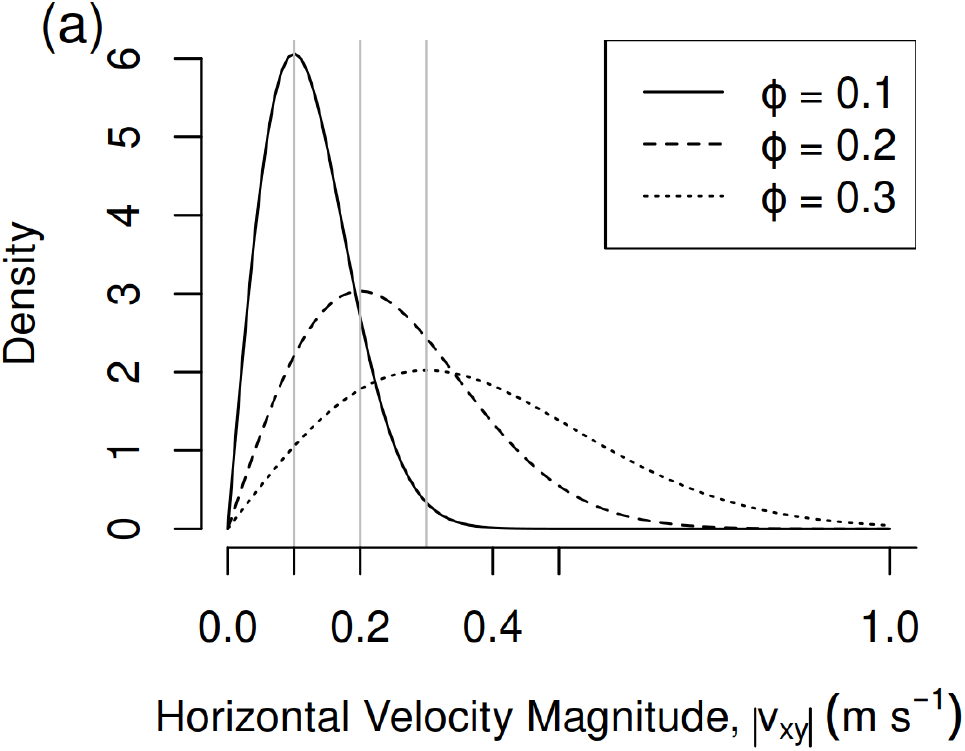
A visualization of the CTCRW velocity process: The stationary velocity magnitudes in the horizontal plane for different values of *ϕ*. Note that the velocity magnitudes follow a Rayleigh distribution with a mode of *ϕ*.

*ϕ* and *t*_05_ can be chosen *a priori* to specify a stationary velocity distribution and correlation structure which is assumed to resemble the animal’s movement in its most active state. Additionally, when fitting EK-TOA, the suitability of the chosen parameters can be checked by examining the distribution of pre-fit time-difference-of-arrival residuals. A visual guide on how to generate these residuals is provided in the supplementary material.

Lastly, forcing a small amount of correlation into the positioning model can reduce stationary jitter — an artefact where detection error is incorrectly attributed as animal movement, resulting in a stationary animal being observed as having large erratic steps centred around a fixed point.

### 3.2 Simulated study

EK-TOA was verified using a time-of-arrival dataset taken from simulated movement tracks. The tracks were generated within a real telemetry array configuration which had previously been used for long-term tracking of two fish species, common barbel (*Barbus barbus*) and European grayling (*Thymallus thymallus*), at the Altusried hydropower plant on the River Iller, south Germany (see Elings et al. 2023; Mawer et al. 2023). The study array was comprised of 16 180 kHz HR2 Innovasea receivers. The water depth has a maximum of 6.9 m and mean of 2.5 m across different discharges and flow velocities averaged 0.34 m s^*−*1^ going up to 1.54 m s^*−*1^ (at a discharge of 40 m^3^ s^*−*1^, the mean annual flow). A satellite image of the study area is provided in Figure 2.

**Figure 2.**
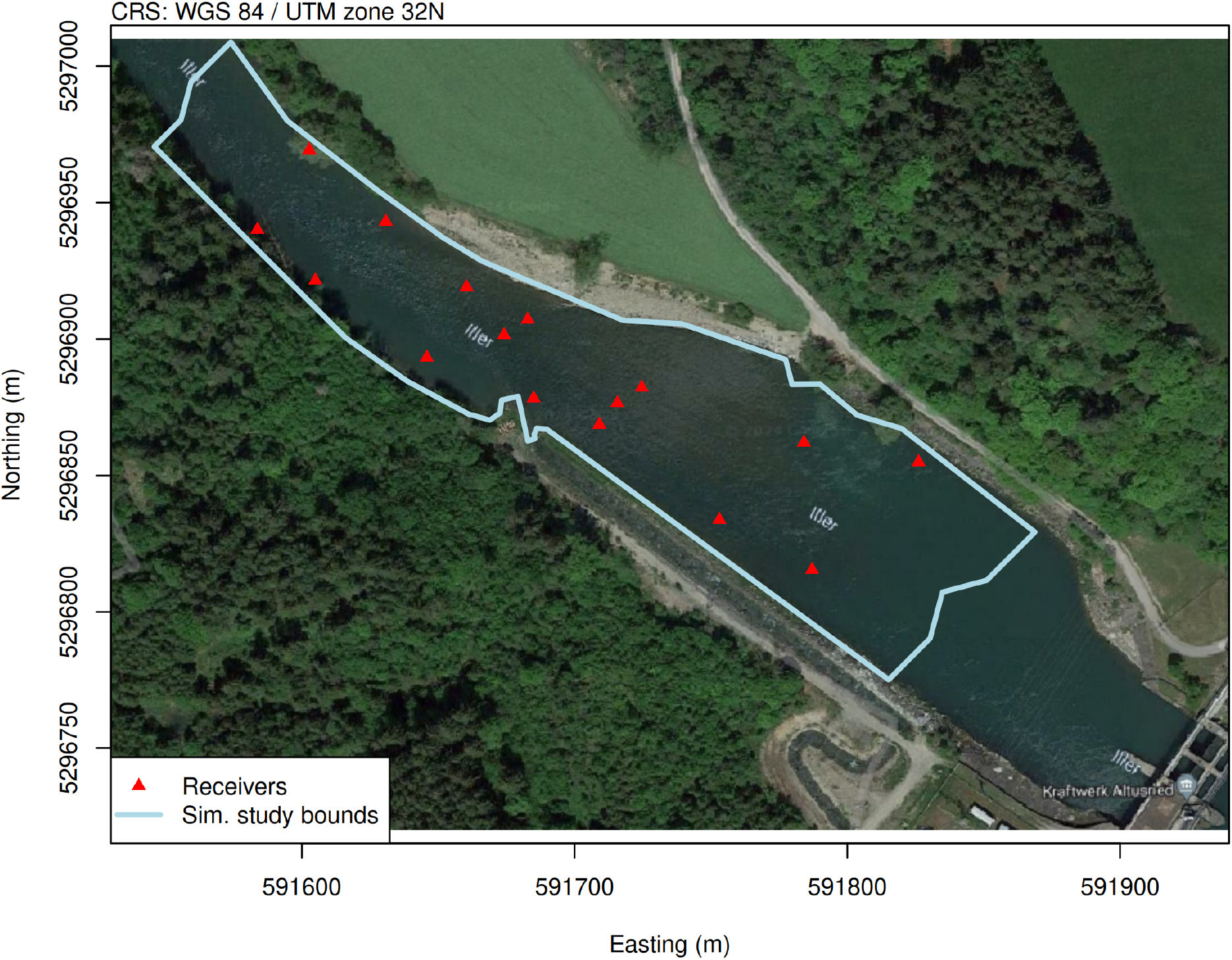
A satellite image of the study area used by both the simulated and real case studies. Receiver locations are plotted along with the bounded area used to generate tracks in the simulated study. The Altusried hydropower plant, shown here, lies on the River Iller in south Germany. Attribution: Imagery © 2024 Maxar Technologies, map data © 2024 GeoBasis-DE/BKG (© 2009).

Johnson et al.’s (2008) CTCRW model was used to create 100 tracks each with 500 points. Each track had a random starting location within the study area which was defined as the stretch of river spanning 50 m up- and down-stream of the telemetry array. A constant emission interval of 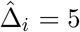 s was used and the movement parameters were set to *ϕ* = 0.2 m s^*−*1^ and *t*_05_ = 100 s. When a track position fell out of the study bounds, the position was relocated to the nearest boundary and the velocity vector for that point was projected to lie parallel to the study bounds, rather than outside it. This resulted in track segments heading out-of-bounds to ‘slide along’ the edge of the study area until the velocity variance directed the track back inwards. Figure 3 shows the simulated CTCRW process as well as a resulting example track.

**Figure 3.**
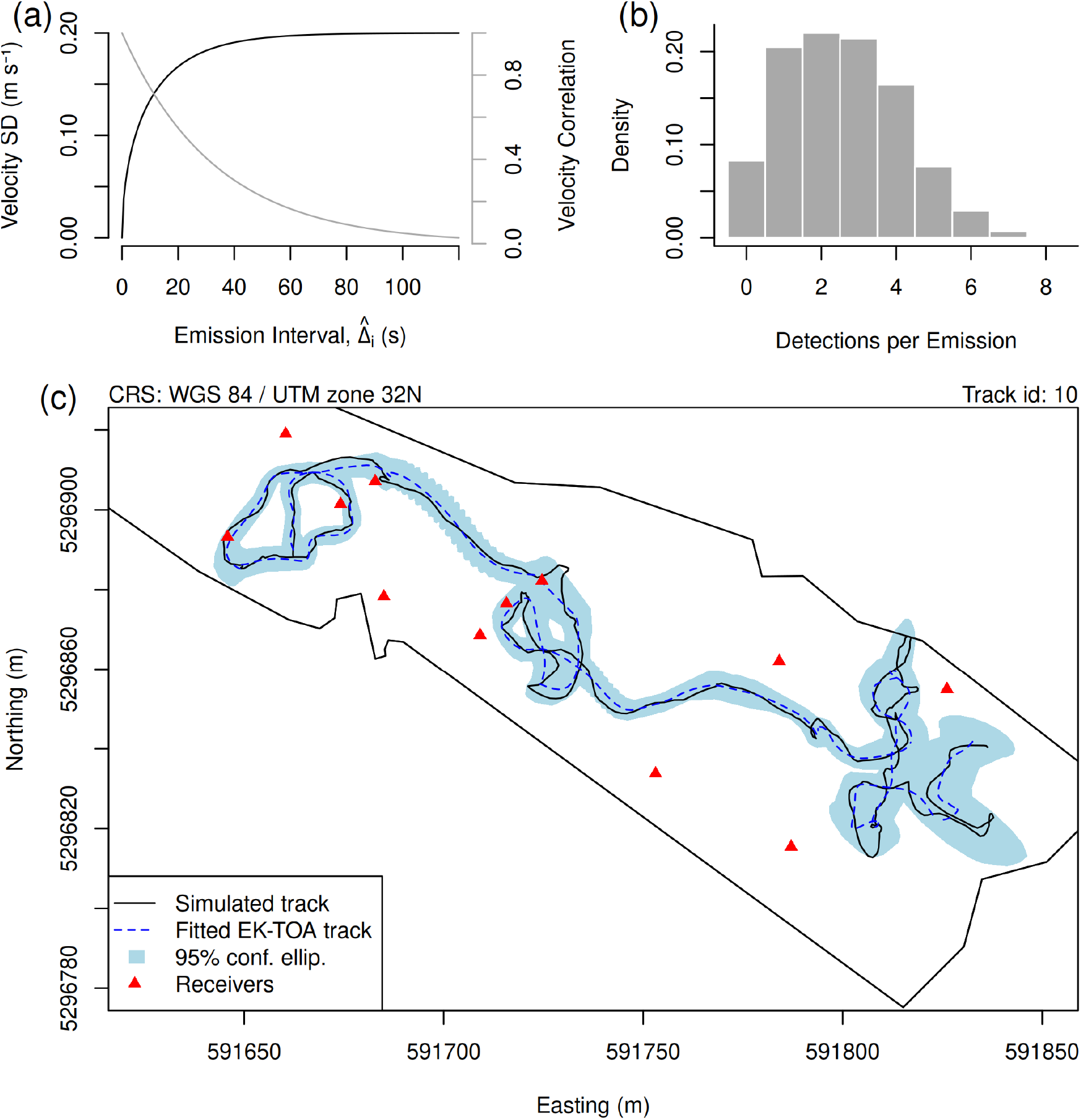
A visualization of the simulated study methods. Panel (a) shows the standard deviation and correlation of the simulated CTCRW velocity process with respect to the length of the emission interval. Panel (b) shows the number of detections per emission, pooled over all simulated tracks. Panel (c) shows an example simulated track along with the estimated track resulting from fitting EK-TOA with the same movement parameters used by the simulated process. The 95% confidence ellipses for all EK-TOA position estimates are shown.

Detection times were simulated by calculating the time-of-arrival for a transmission velocity of 1500 m s^*−*1^ and adding a Gaussian detection error with *σ*_det_ = 2 ms. For all receivers with line-of-sight and within 50 m of an emission, a 0.7 probability of detection was given. Receivers beyond that range or with no line-of-sight were assigned missed detections. These detection parameters were broadly based on previous experience and represent conditions that may resemble a relatively challenging telemetry setup.

Regions of the study area near the edges of the up- or down-stream bounds suffered from low detectability due to far distances from most of the receivers. Tracks in these regions could consist solely of emissions with fewer than 3 detections, which is can give very poor positioning. As a result, the first and last 4 emissions in each track were forced to have at least 4 detections. When there were fewer than 4 detections for these emissions, nearby undetected receivers were changed to detected. Figure 3 reports the resulting detections per emission pooled across all the simulated tracks.

For the time-of-arrival values from each whole track, EK-TOA was fitted using an extended Kalman filter. For emissions with less than two detections, the update step of the Kalman filter was skipped, as these emissions effectively held no positioning information. After running the filter, state estimates were further refined using a Rauch–Tung–Striebel (RTS) smoother (Rauch et al. 1965). The RTS algorithm updates the posterior estimates for the mean and variance of each state using all measurements in the time-series — as opposed to only past measurements, as done by the filter. While there are multiple smoothing algorithms to choose from, the RTS algorithm was chosen here due to its ease of implementation. Additionally, since the process model of EK-TOA is linear, the RTS algorithm can be applied here in the same manner as in a Kalman filter.

Extended Kalman filters require an initial estimate for the first state to be provided along with a covariance matrix for that estimate. This was set agnostically where the first state position was set to the center of the study area with a standard deviation equal to the distance between this central point and the farthest receiver. This effectively tells the filter that the first emission occurred somewhere in the general vicinity of the array. The first velocity and emission error states were set to 0; first velocity variance to *ϕ*^2^; and the first emission time error variance to 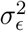.

Multiple EK-TOA models with varying movement parameters which both over- and under-estimated the simulated process were fit to the dataset. The movement parameters were set to unique combinations of *ϕ* = 0.05 m s^*−*1^, 0.1 m s^*−*1^, 0.2 m s^*−*1^, 0.4 m s^*−*1^ and 0.6 m s^*−*1^ and *t*_05_ = 1 s, 5 s, 10 s, 25 s, 100 s and 400 s. All models had an observation error, *σ*_det_, set to the same value used to create the simulated detections. The emission error standard deviation was set to *σ*_*ϵ*_ = 0.33 s following the protocol suggested in the previous section.

After fitting the models, absolute positioning error was measured by taking the distances from the true simulated positions to the paired EK-TOA estimates. Additionally, standardized positioning error values were generated to examine how well the distribution of absolute positioning error values matched with the positioning error covariance returned from the model. For each emission, an error vector was taken by subtracting the EK-TOA position estimate from the true simulated position. Next, Mahalanobis distances (Mahalanobis 1936) of these vectors were calculated using the positioning error covariance returned by the EK-TOA model. If the absolute positioning error matches well with the estimated positioning error covariance, the resulting standardized positioning error is expected to fall along a *χ*_2_ distribution (a chi distribution with two degrees of freedom). Standardized positioning error that deviates from a *χ*_2_ indicates the EK-TOA model is over- or under-estimating the absolute positioning error.

The positioning error covariance returned by EK-TOA can be taken as the model’s estimate of its own positioning accuracy. If this is well fitting — as measured by the standardized positioning error — the 95% confidence ellipses for EK-TOA positions can be interpreted as reasonably accurate.

Finally, to compare how well the standardized positioning errors matched with the expected *χ*_2_ distribution across EK-TOA models with different movement parameters, one-sample Kolmogornov-Smirnov (KS) statistics were calculated for each fit. Ranging from 0 to 1, lower values indicate the standardized positioning error distribution more closely matches with the expected *χ*_2_ distribution. Hence, a lower KS value for a fit suggests that the 95% confidence ellipses for those position estimates are more trustworthy.

### 3.3 Real study

To provide a real-world example of EK-TOA, it was fitted to a real dataset collected from the site described in the simulation study. Many fish (22 common barbel, 6 European chub, and 25 European grayling) in this site were tracked using Innovasea V9-180 acoustic tags surgically implanted in the peritoneal cavity. The tags were configured to simultaneously emit both *pulse position modulation* (PPM) and *high residency* (HR) transmissions — the proprietary communication protocols supported by Innovasea receivers — with random emission intervals ranging from 50 s to 70 s and 1.1 s to 1.3 s, respectively.

A 24h period from this dataset was selected for the case study on the date 2018-08-23. Here, the receiver clocks in the array were synchronized with a hierarchical model coded in the TMB package in R. Likely reflected transmissions were then removed with a population Monte Carlo algorithm that provided the probability that a given detection resulted from a reflected transmission (Campbell et al. 2025). Those giving a probability larger than 0.5 were removed from the time-of-arrival dataset. The resulting dataset consisted of a combination of both PPM and HR detections.

From this 24 h dataset, a single tagged individual was chosen for the case study which (i) spent most of its time within the array bounds, and (ii) also resulted in a large number of detections. The time-of-arrival data from the selected individual resulted in approximately 17 h of tracking data comprised of 28244 detections from 5403 emissions. The chosen individual was a common barbel caught on 2018-05-17 during its upstream spawning migration.

EK-TOA was fitted to the case study time-of-arrival data using the same protocol from the simulated study methods. The detection and emission parameters were kept the same as the simulated study. Upon examining the detection error distributions of the sentinel tags in the study area, the measured *σ*_det_ of the dataset was determined to be nearer to 1 ms. However, the value used by the EK-TOA was left at 2 ms to provide more cautious position estimates, as the observed detection error distribution was heavily tailed. A description of the sentinel tag error is provided in the supplementary materials.

The movement parameters for EK-TOA were selected by examining the resulting standardized pre-fit difference residuals (see supplementary material), giving *ϕ* = 0.1 m s^*−*1^ and *t*_05_ = 5 s. These chosen parameters were selected to avoid overdispersion (see Figure 7), as our simulated analysis showed overdispersed residuals tended to result in poor measures of positioning uncertainty.

After fitting, a 95% confidence polygon was drawn around all the estimated positions. For each state position, the Cholesky decomposition of the covariance matrix was used to generate a 95% confidence ellipse. So long as the absolute error distribution is approximated well by the EK-TOA model, 95% of the true tag positions are expected to lie within these ellipses. Across all states, these confidence ellipses were merged into a single polygon and plotted. The resulting polygon gives a simple visual indicator of the positioning uncertainty along the EK-TOA track.

Additionally, positioning accuracy measures were calculated from the 95% confidence ellipses. For each position estimate, the distance was measured from the center of the confidence ellipse to its farthest edge — the semi-major axis. These values provide a rough estimate of the upper limit of the expected absolute positioning error for a given emission.

For comparison, TDOA positioning was also done on the prepared time-of-arrival dataset. For all emissions which had 4 or more detections, Smith and Abel’s (1987) spherical-interpolation method was used to estimate tag positions. This method gives an approximate closed-form solution to TDOA positions for 2D arrays consisting of 4 of more receivers. While there are numerous methods to implement TDOA positioning, the spherical-interpolation method is fast, simple to implement, and returns positions with a comparable accuracy to those provided by popular commercial telemetry positioning services.

## 4 Results

In the simulated study, EK-TOA models which used the same movement parameters as the simulated process resulted in good fits. The majority of the absolute positioning error fell below 5 m, and the standardized positioning error closely matched the expected *χ*_2_ distribution (Figure 4), indicating the estimates of positioning uncertainty were reliable.

**Figure 4.**
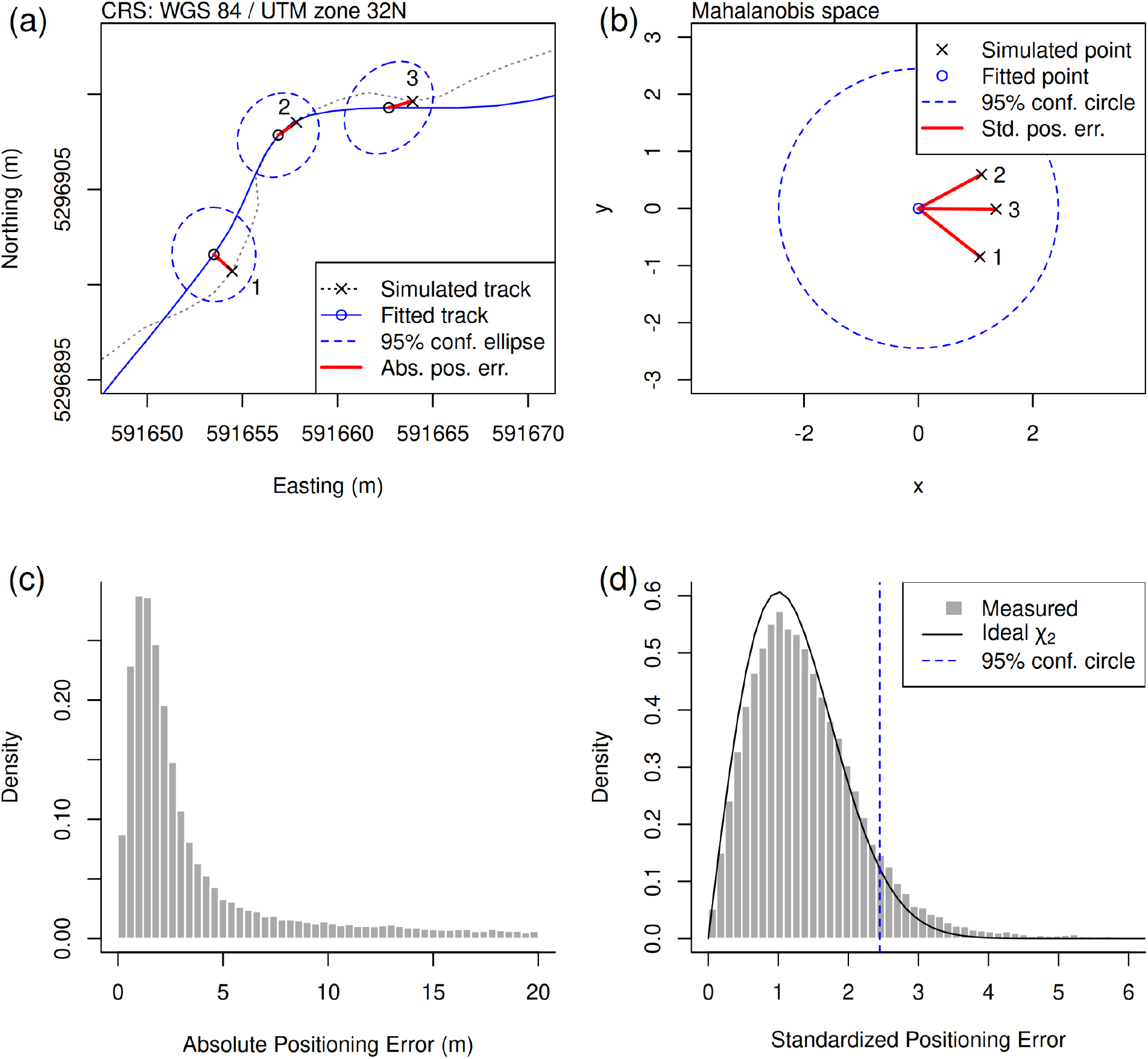
Results from fitting EK-TOA to the time-of-arrival data generated from 100 simulated tracks. Here, EK-TOA was fitted using the same movement parameters that were used to generate the simulated dataset. Panel (a) shows a section of a simulated track along with the fitted EK-TOA track. Three points are chosen to illustrate the absolute positioning error. Panel (b) shows those same three points in Mahalanobis space. Here, the absolute positioning errors are standardized by their respective 95% confidence ellipses, giving the standardized positioning error. Panels (c) and (d) give the absolute and standardized positioning errors pooled across all 100 tracks. When the positioning uncertainty returned by the EK-TOA model — confidence ellipses — matches well with the absolute positioning error, the standardized error will fall along a *χ*_2_ distribution. Here, the EK-TOA model slightly underestimated the absolute positioning error.

When examining the EK-TOA models fitted with variable movement parameters (Figure 5), simultaneously over- or under-estimating both *ϕ* and *t*_05_ resulted in large increases in positioning error. Notably, combinations of overestimating *ϕ* while underestimating *t*_05_, or vice versa, resulted in only small increases in positioning error, reflecting the dependence structure between these two parameters in the resulting error. For example, the parameter combinations of *{ϕ* = 0.05 m s^*−*1^; *t*_05_ = 400 s*}* and *{ϕ* = 0.6 m s^*−*1^; *t*_05_ = 5 s*}* both resulted in a mean positioning error distribution similar to selecting the true simulation parameters of *{ϕ* = 0.2 m s^*−*1^; *t*_05_ = 100 s*}*.

**Figure 5.**
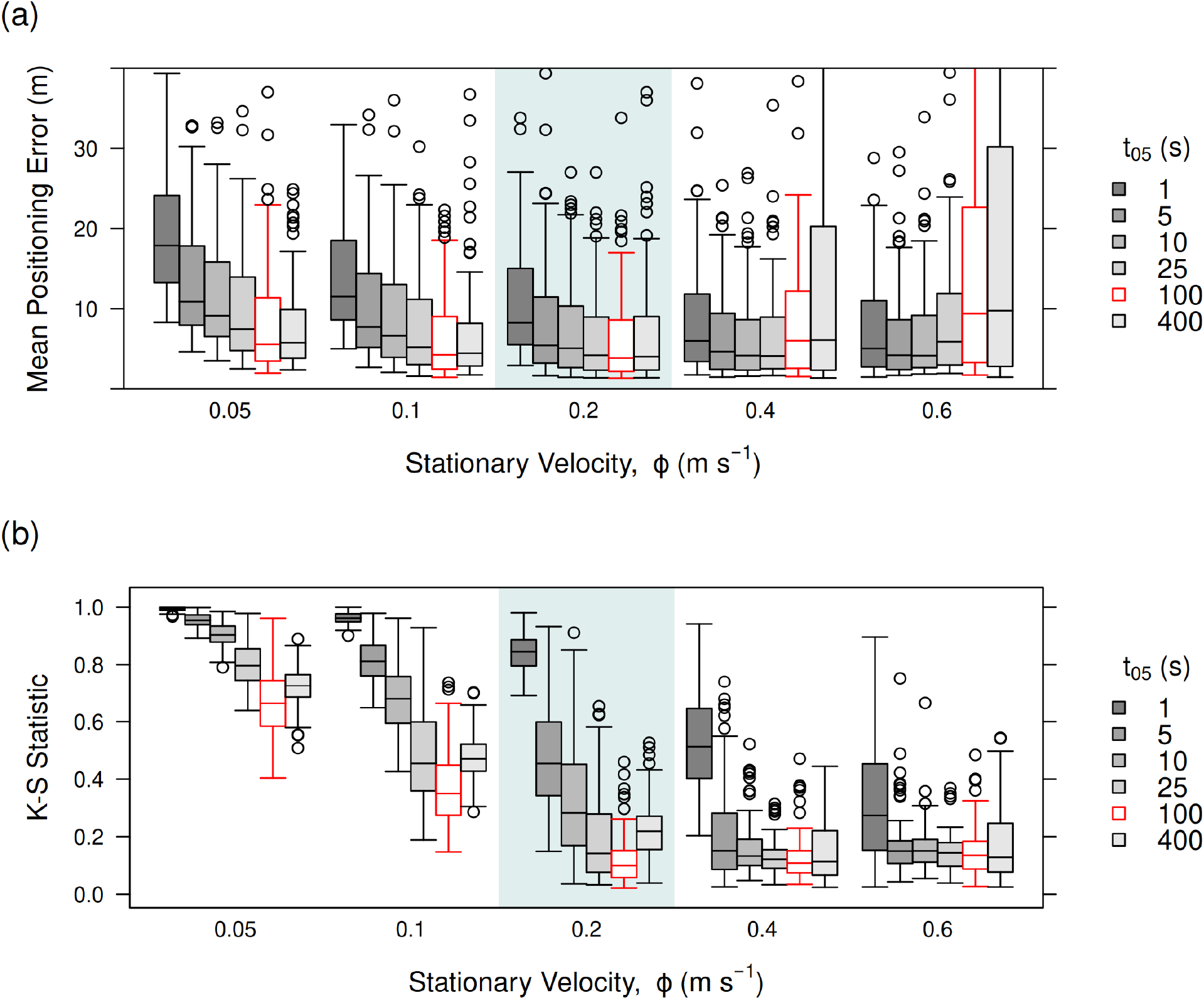
Results from fitting EK-TOA models with variable movement parameters to the 100 simulated tracks. The movement parameters used to fit the EK-TOA models are shown in the x axes and boxplot legends. Those values used by the simulated process are marked with a grey section along the x axes and red outline around the boxplots. Panel (a) shows the mean positioning error returned by each EK-TOA fit. Panel (b) shows the Kolmogornov-Smirnov statistic for each fit, comparing the standardized positioning error to an ideal *χ*_2_ distribution. Lower values indicate more accurate estimates of positioning uncertainty.

Trends in the standardized positioning error, however, were different. Here, underestimating *ϕ* resulted error distributions that deviated farther from the expected *χ*_2_ distribution, as evidenced by the increasing KS values. Other combinations of parameters had relatively small effects on the KS values.

In the supplementary material, distributions for the standardized pre-fit difference residuals are reported for these EK-TOA fits. It is interesting to note that fits resulting in high KS values (i.e. poorly estimated positioning uncertainty) were correlated with residual distributions that were overdispersed. Conversely, residual distributions that were underdispersed (*{ϕ* = 0.6 m s^*−*1^; *t*_05_ = 25 s*}*, for example) still provided relatively low KS scores.

Finally, both EK-TOA and TDOA positioning were applied to the real case study dataset (Figure 6 & 7). Here, EK-TOA gave extremely small positioning accuracy values, largely falling below 4 m. Upon examining the tracks, with one exception, the entire track fell within the known bounds of the river. Note that as the start/end segments of tracks have less data to make use of, they are prone to higher error than the intermediate sections of the track. This is an inherent feature of Markovian animal movement models, including YAPS. As the track moved to the up- and down-stream edges of the study area, positioning accuracy decreased — as seen by larger 95% confidence ellipses for those positions. The TDOA position estimates, on the other hand, were highly variable, frequently giving position estimates which fell well outside the known study area bounds.

**Figure 6.**
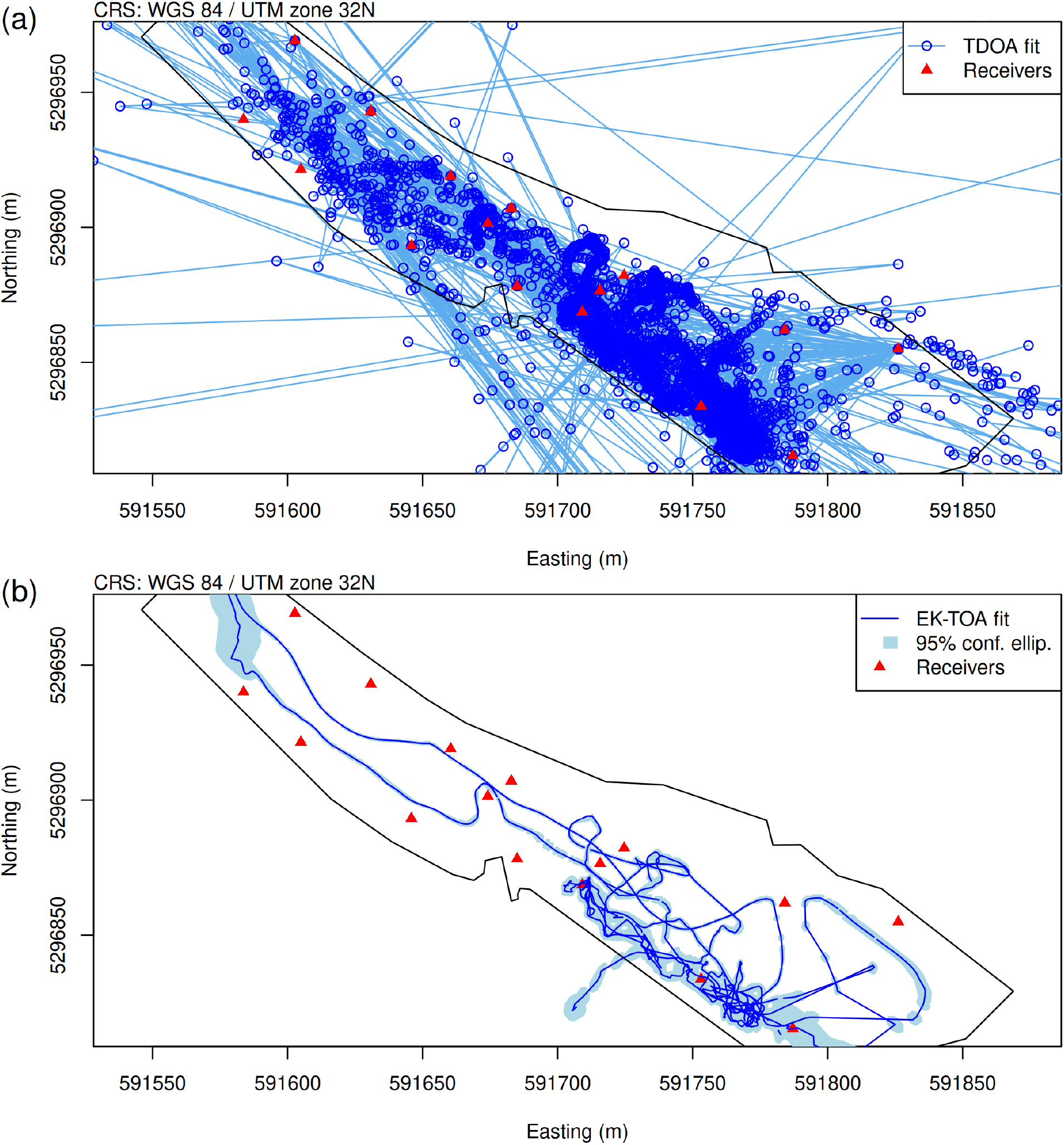
Real case study where the EK-TOA model was fitted to 17 hours of continuous telemetry data from a common barbel. Panel (a) shows position estimates returned by TDOA positioning. Panel (b) is the EK-TOA estimates. The 95% confidence ellipses for EK-TOA position estimates give a visual indicator of positioning uncertainty.

The residual diagnostic plots are shown in Figure 7, panels (c) and (d). Here, the residuals were underdispersed, where the observed values fell well within the bounds of our expected residual distribution. From our simulated study, it can be expected that such underdispersion increases in the absolute positioning error while still providing good values of positioning uncertainty.

**Figure 7.**
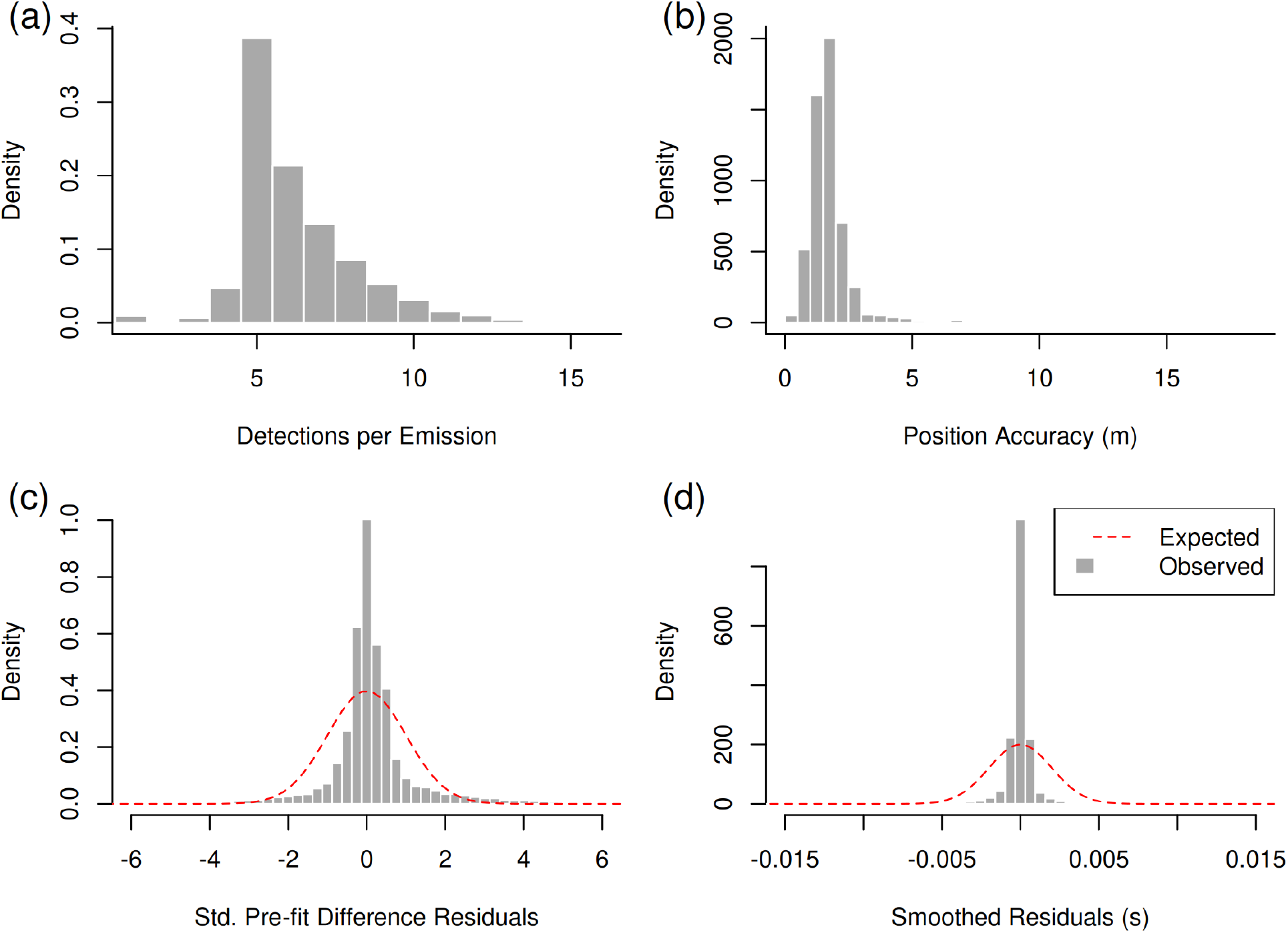
EK-TOA diagnostic plots from the real case study. Panel (a) shows a histogram of the number of detections per emission event. Panel (b) gives the positioning accuracy; the length of the semi-major axis from the 95% confidence ellipses for each estimated position. Panel (c) shows standardized time-difference-of-arrival residuals returned from the predict step of EK-TOA. Panel (d) gives the time-of-arrival residuals returned after smoothing has been applied to the state estimates.

Finally, the estimated instantaneous velocity values are provided in the supplementary material. Here it can be seen that the tagged animal has brief bursts of high velocity travel, while for the remainder of the track it expresses low velocity, inactive movement behaviours.

## 5 Discussion

The EK-TOA model presented here allows for positioning large-volume telemetry datasets that may otherwise require complex processing pipelines or large computational resources when using state-space models fitted by iterative optimization procedures, such as YAPS. The results from the simulated study verified that EK-TOA can give positioning estimates with both (i) small absolute positioning error and (ii) well fitting error distributions. In addition to high accuracy, the EK-TOA model is extremely fast to execute. In the example real case study, the model required less than 1 second to fit the more than 5000 positions (with an Intel i5–13600K CPU, written in R).

As EK-TOA position estimates are closed-form, the processing time for a dataset with a given distribution of receiver detections per tag emission is expected to scale linearly with the volume of data, irrespective of the number of tagged individuals, positions of the tags, tag-emission intervals, and geometry of the receiver array setup. In the supplementary material, a pure R implementation of the model is provided that has been written for the purposes of readability over optimization. For even faster processing, EK-TOA can be coded more efficiently within R or implemented in compiled and just-in-time compiled languages such as c/c++ and Julia.

The simulated time-of-arrival dataset was designed to approximate the detection error observed in real telemetry arrays (a Gaussian distribution with *σ*_det_ = 2 ms). In practice, detection error can have non-normal kurtosis and skew in addition to other spatially dependant error properties. As well as the inherent detection time error on telemetry receivers, clock drift between independent receivers, the properties of the acoustic propagation environment, and uncertainty in the positions of the telemetry receivers can add additional sources of error which are difficult to capture with a Gaussian. A key limitation of the EK-TOA model is the requirement that the detection error distribution must be Gaussian — a constraint of the extended Kalman filter. To capture heavily tailed detection error, EK-TOA will require specifying a sufficiently broad Gaussian distribution.

In addition to the above mentioned sources of detection error, reflected transmissions can cause late-arriving detection error outliers that are orders of magnitude greater than the those from direct transmissions. In environments with acoustically reflective surfaces, such as dams, weirs, or fish passes, these reflected tag transmissions can prevent the application of any telemetry positioning model if they are not handled. In the current case study, large reflected transmissions were identified and removed before the application of the EK-TOA and TDOA models using the PMC-TOA model described in Campbell et al. (2025).

In the case that a small number of large detection outliers are present when fitting the EKTOA model, their identification and removal can be straightforward. The extended Kalman filter provides position estimates in a time-ordered sequence. For a dataset with a single large detection outlier in the middle of a track, the position estimates before the outlier will resemble a well fitted track. At the outlier, the track will either have a transient position outlier at that point or the entire latter segment of the track will be become unstable and provide clearly recognizable incorrect position estimates — such as being far outside the array or having extremely large steps. Here, one can simply remove the emissions near the point where the track became unstable and refit EK-TOA.

State-space models such as EK-TOA and YAPS are useful tools for telemetry positioning but their performance can be difficult to validate. These models achieve good positioning accuracy by making assumptions about the underlying movement process of the tagged animal. As the animals movement behaviour deviates from the model’s assumptions, state-space models may become less accurate. While it may seem reasonable to attach a telemetry tag to a boat to evaluate the performance of a state-space model with the measured GPS tracks, the movement behaviours of a fish are likely to be quite different and more difficult to observe with a state-space model than what can be replicated with a driven boat, particularity at time resolutions on the order of a few seconds between emissions. While ground-truthed GPS data is a useful validation tool, further extrapolation to tagged fish should be done with caution from these datasets, as they likely do not capture the fine-scale complexities and changing behaviour states of real fish. For example, replicating movements similar to random foraging — which may be of interest to researchers and difficult to observe with state-space models — with a GPS tracked tag is practically challenging.

EK-TOA and YAPS are similar in the sense that they are both state-space models with a time-of-arrival observation model and a random-walk process model. Beyond the different fitting procedures, YAPS models the tag emission process (static or random emission intervals) to predict tag emission times while EK-TOA treats emission times as an error term of some initially provided guess. The former approach restricts datasets to be comprised of a single tag type with a known emission process. The latter requires no information about the tag emission processes and allows fitting datasets comprised of composite tags, such as the combination PPM & HR emission protocol used in the case study.

Additionally, YAPS can fit non-Gaussian time-of-arrival errors (such as a mixture of t and Gaussian) while EK-TOA cannot. While selecting heavier-tailed detection error distributions can aid positioning in the presence of large reflection outliers (see Vergeynst et al. 2020), reflected detections provide no positioning information and can make the fitting procedure more computationally challenging and prone to failure. Hence, the removal of large reflection outliers during preprocessing, as done here, provides a benefit to both YAPS and EK-TOA. Beyond the consideration of reflected detections, the flexibility of selecting non-Gaussian error distributions can allow for more accurate position estimates.

With respect to the process models, EK-TOA applies a CTCRW movement model while YAPS applies a Gaussian random walk. When the CTCRW relaxation parameter, *β*, is set high, the CTCRW approaches a Gaussian random walk, as in YAPS. When set lower, the added ‘momentum’ in the estimated track can aid in reducing positioning error. Here, the CTCRW provides a more suitable movement model for fine temporal resolution data where correlated movements are expected. Lastly, a Gaussian random walk assumes a linear relationship between the time interval and the variance of the animal’s displacement. At short intervals, where correlation is present, the CTCRW models this relationship as non-linear. However, over larger time steps, this relationship converges towards a linear trend that approximates *α*^2^ *· β*^*−*2^. This can be a useful measure to compare the displacement processes resulting from a Gaussian random walk and a CTCRW.

YAPS uses maximum likelihood to fit the movement and observation parameters while the EKTOA workflow presented here set these parameters *a priori* and checked their suitability by viewing the standardized pre-fit difference residuals. In the case of the simulated study, erring on the side of overestimating the variance of the true velocity process, *ϕ*, prioritizes good measures of positioning uncertainty over positioning accuracy. This may result in larger absolute positioning error, however, the resulting 95% confidence ellipses tend to be more accurate than if *ϕ* is under-estimated.

It may be desirable to fit one or both of the EK-TOA movement parameters via maximum likelihood, and this is simple to to with the current model. In this case, careful thought should be given to how the fitted movement parameters affect the resulting estimated track, particularity when making behavioural inference from comparing tracks fitted with different movement parameters. Even when using optimally fitted parameters, it can still be informative to check the standardized pre-fit difference residuals as a diagnostic measure, as this will inform the interpretation of the resulting measures of positioning uncertainty.

The extended Kalman filter utilized by EK-TOA is just one particular type of recursive filter. Future work on telemetry positioning can explore the use of more flexible recursive filters where non-Gaussian error distributions can be fitted (see Särkkä and Svensson 2023). Recursive filters which allow for complex detection error distributions can potentially remove the need for the pre-processing step of removing large outliers resulting from reflected detections while still maintaining fast processing times. For example, particle filters have recently been shown to be a promising tool for tracking aquatic animals through acoustic telemetry networks (Lavender et al. 2023; Lavender et al. 2024).

Note that telemetry setups have unique patterns of spatial positioning error and detectability which can vary over time and every study area should be treated as a unique case. Hence, the simulated and case studies presented here should be interpreted with caution when generalizing to other setups. For the application of any telemetry positioning model to a dataset, it is recommended to compare candidate positioning models using simulated datasets, as done here, to better understand (i) how readily movement behaviour can be disentangled from detection error and (ii) how parameter selection impacts the resulting positioning error.

Additionally, within the telemetry community there is a need for a curated set of benchmark datasets which can be used to compare the output and computational costs of available positioning models over a broad set of study areas (Campbell et al. 2025). With such a dataset, researchers would be able to set realistic expectations on how each positioning model may perform on their particular study area.

As the temporal resolution of acoustic telemetry tags increases and deploying large, dense receiver arrays becomes more feasible, researchers are becoming increasingly interested in examining animal behavioural processes on these fine scales. For studies examining switching between behavioural states and the navigation cues driving each of these states, hidden Markov models (HMMs) and step selection functions (SFFs) can provide the respective statistical inference (see Elings et al. 2023; Mawer et al. 2023). These types of behavioural analyses are simplified by treating the data as having discrete time units. As acoustic telemetry signals are often emitted in randomized time intervals, telemetry positions may need to be interpolated to match the desired discrete time scale. For EK-TOA, interpolation is simple as it is an intrinsic feature of Kalman filters. By providing 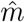 values for the desired times with no detection data, EK-TOA will return the interpolated positions and covariances.

In conclusion, the EK-TOA model presented here is shown to be a capable positioning model for large telemetry datasets. For practical advice on applying EK-TOA, the following rough guidelines are given: (i) ensure there are no large outliers resulting from reflected detections in the time-of-arrival dataset; (ii) utilize sentinel tags to empirically set a suitable detection error distribution; (iii) apply EK-TOA with either (a) CTCRW parameters set *a priori*, based on the animal’s expected behaviour in its most active state, or (b) fit them using maximum likelihood; (iv) after the model has run, make use of the standardized pre-fit difference residuals as a diagnostic tool to aid in the interpretation of the measures of positioning uncertainty; (v) when initially applying the EK-TOA, do not apply smoothing so that problematic emissions with large detection outliers can be more easily identified and removed. Finally, R implementations of the EK-TOA model, the example case study dataset, and the functions for generating simulated data have been provided in the supplementary materials along with reproducible examples of fitting the model.

## Supporting information

Example implementation in R

Detection Error

Pre-fit Residuals

Real case study figures

Example code and data

## Acknowledgements

This project has received funding from the European Union Horizon 2020 Research and Innovation Programme under the Marie Sklodowska-Curie Actions, Grant Agreement No. 860800.

The data presented in the real study was collected as part of the FITHydro project, a European Union’s Horizon 2020 (H2020) research and innovation program under Grant Agreement No. 727830.

## Author Contributions

JC developed the model, executed the analysis, and wrote the manuscript & supplementary material. JE curated the case study dataset. PL & RM reviewed the supplementary material. IP collected the data used by the case study. All authors reviewed and contributed to the manuscript.

## Declaration of Generative AI and AI-assisted technologies in the writing process

Generative AI or AI-assisted technologies were not used at any stage in the preparation of this manuscript or the work presented in the manuscript.

