## Supplementary material for "An extended Kalman filter for large-volume path positioning of aquatic animals within acoustic telemetry arrays": Example implementation in R

This document provides reproducible examples of fitting the EK-TOA model described in the manuscript. All example data and required code is found in the supplied `./data` and `./code` directories, respectively. For all custom functions, brief documentation can be found in the comments above the function definitions within the `./code` directory.

```
# Load required packages
require(sf)      # GIS
require(units)   # GIS
require(chi)     # For chi distribution functions
require(mvtnorm) # For simulating data
```

### Fitting the EK-TOA model

Here, the case study results from the manuscript (Figure 6) will be reproduced. First, load the case study dataset.

```
# --- Define study site CRS
crs <- st_crs("EPSG:32632")
# --- Load receiver array (matrix)
f_rec <- "./data/receiver_locations.csv"
mat_rec <- read.csv(f_rec, row.names = 1) > as.matrix()
pts_rec <- st_sfc(st_multipoint(mat_rec), crs = crs)
# --- Load time-of-arrival dataset of tagged fish (matrix)
f_toa <- "./data/toa_mat.csv"
mat_toa <- read.csv(f_toa) > as.matrix()
colnames(mat_toa) <- rownames(mat_rec)
# --- Load polygons of study area (SFC_POLYGON)
f_site <- "./data/study_site_full.csv"
ply_site <- read.csv(f_site) > as.matrix() > list() >
  st_polygon() > st_sfc(crs = crs)
f_site_clip <- "./data/study_site_clip.csv"
ply_clip <- read.csv(f_site_clip) > as.matrix() > list() >
  st_polygon() > st_sfc(crs = crs)
```

Make sure the receiver order in the columns of the TOA matrix match the rows of the receiver array matrix. NA values indicate missed detections and all units are in meters and seconds. For emissions that result in less than 2 detections, EK-TOA will skip the update step for these states.

Here's a labeled overview of the receiver array.

```
plot(ply_clip, axes = T,
     xlab = "Easting (m)", ylab = "Northing (m)", border = 'darkgray')
plot(pts_rec, pch = 17, col = 'red', add = T)
mtext("CRS: WGS84, UTM Zone 32N", adj = 0)
```

```
text(labels = rownames(mat_rec),
      x = mat_rec, adj = c(0,0), col = 'black', cex = 1)
```

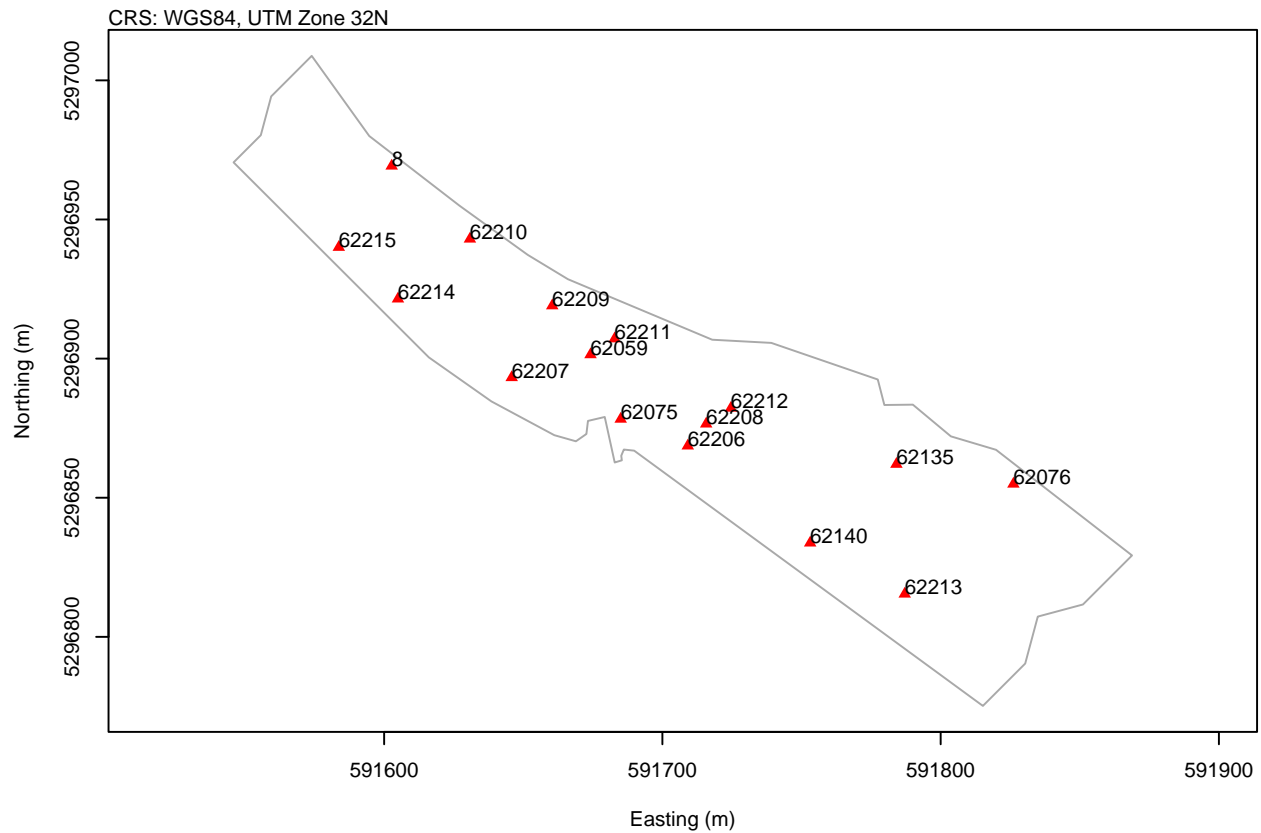

The functions to apply EK-TOA can be loaded from `./code/EK-TOA_main.R`. First, set the continuous-time correlated random walk (CTCRW) movement model parameters to some sensible *a priori* values. For this case study, $\phi = 0.15 \text{ m s}^{-1}$ and $t_{05} = 15 \text{ s}$.

```
# --- Mode of the uncorrelated velocity magnitude distribution (see fig. 1)
phi <- 0.15# (m/s)
# --- Time interval where the velocity correlation falls below 0.05
t_05 <- 15# (s)
# --- Convert to CTCRW parameters from Johnson et al. (2008)
# Equations (6) & (7) from the manuscript
beta <- 3/t_05
alpha <- phi*sqrt(2*beta)
```

Next, set the initial state for the Kalman filter to start on. An agnostic choice is a position in the center of the array with a standard deviation matching the distance to the farthest receiver. This effectively tells the Kalman filter the first emission occurred somewhere in the general vicinity of the array.

```
# --- Get central point inside array
pt_init_pos <- st_convex_hull(pts_rec) > st_centroid()
init_pos <- st_coordinates(pt_init_pos); init_pos

##           X           Y
## [1,] 591704.3 5296890

# --- Distance from initial position to farthest receiver
init_pos_sd <- st_distance(pt_init_pos, st_cast(pts_rec, to = "POINT")) >
```

```
max() > as.numeric()); init_pos_sd
```

```
## [1] 130.4658
```

The code block below will run EK-TOA. Here, the estimated emission times, $\hat{m}_i$, are automatically generated. The values for the first velocity and emission error state are also set automatically. The argument `smooth = T`, will apply a Rauch-Tung-Striebel smoother to the resulting positions. Disabling the smoother can be useful for locating detections outliers if the extended Kalman filter becomes unstable — this can be caused by extremely large detection outliers, such as those from reflected detections.

```
# --- Load EK-TOA model
source("../code/EK-TOA_main.R")
# --- Detection error standard deviation
sigma_det = 2e-3# (s)
# --- Speed of signal transmission
c = 1500# (m/s)
# --- Maximum propagation distance
# This is used to automatically set a value for sigma_{emis}
d_max = 500# (m)
# --- Fit the model
ektoa <- eKalTOA(
  toa_matrix = mat_toa, r = mat_rec, c = c, d_max = d_max, # Data
  init_s = init_pos, init_s_sd = init_pos_sd, # Initial state
  sigma_det = sigma_det, # TOA model parameters.
  alpha = alpha, beta = beta, # CTCRW model parameters.
  smooth = T # Enable the RTS smoother after fitting.
)
# --- Show summary of model fit
ektoa
```

```
## Fitted EK-TOA model
##
## --- Time-of-arrival summary
## - Emissions: 5403
## - Median emission interval: 1.78 s
##
##      Number of detections per emission
##      0 1 2 3 4 5 6 7 8 9 10 11 12 13 14 15
## Count 51 5 33 255 2091 1154 726 461 284 166 84 54 22 12 4 1
##
## --- Fixed parameters
## Movement model (continuous-time correlated random walk)
## - Vecolity alpha.: 0.094868 m s<sup>{-3/2}</sup>
## - Velocity beta: 0.200000 s<sup>{-1}</sup>
## - Stationary velocity std.dev: 0.150000 m s<sup>{-1}</sup>
## - Time until corr. falls below 0.05: 15.000000 s
##
## Observation model (time-of-arrival positioning)
## - Detection std.dev.: 0.002000 s
## - Emission error (epsilon) std.dev.: 0.333333 s
```

`eKalTOA()` will return an S3 object for which a number of convenience methods have been written. `print.eKalToa()` will give you an overview of the model parameters and summary information about the input TOA data. The fitted track can be plotted and the fitted position and emission times can be extracted.

```
# --- Get fitted positions
state_position(ektoa) > head()
```

```
##           s.x      s.y
## [1,] 591705.5 5296822
## [2,] 591705.1 5296822
## [3,] 591703.4 5296821
## [4,] 591703.2 5296821
## [5,] 591703.1 5296821
## [6,] 591702.9 5296820
```

```
# --- Get fitted emission times
state_emission(ektoa) > head()
```

```
## [1] -0.2544179  1.9099478 10.1863580 11.3460814 12.5876056 13.7654370
```

```
# --- Show fitted track
plot(ektoa)
```

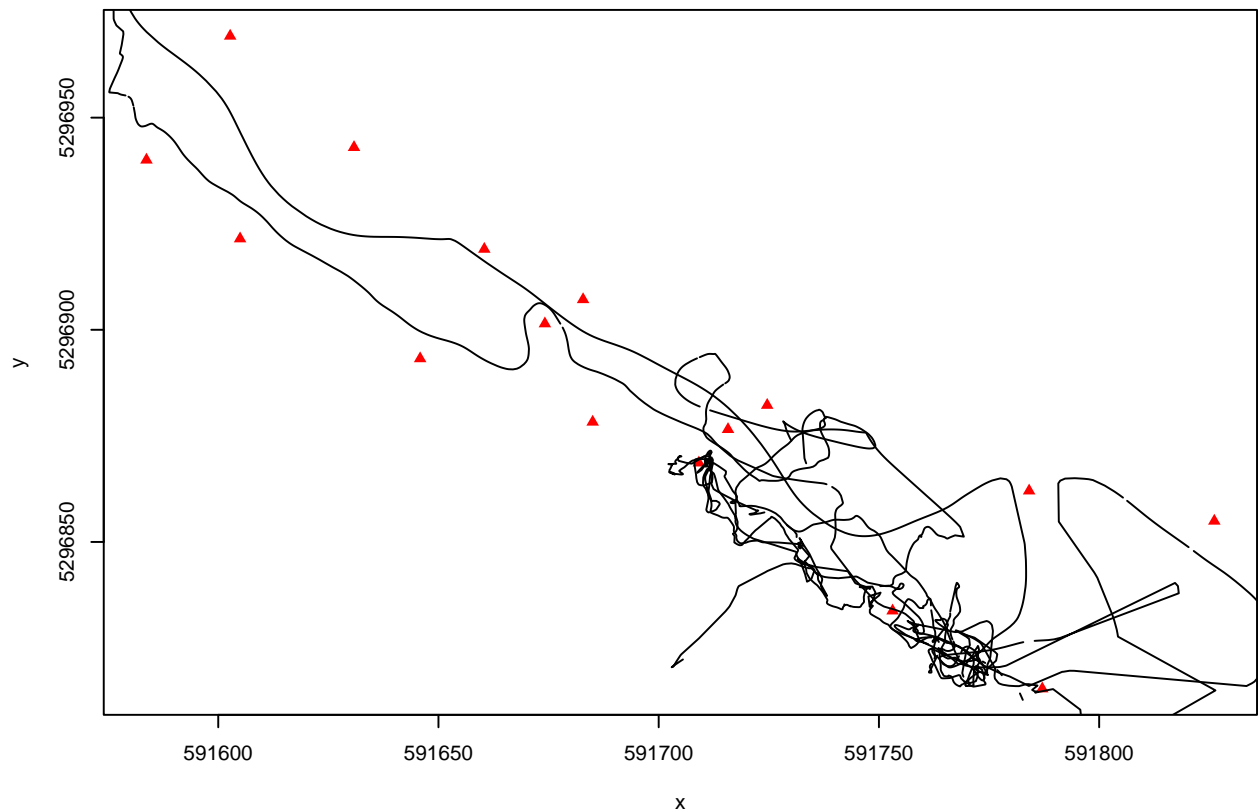

After fitting the model, it's good practice to check your residuals to see if the selected parameters do a reasonable job of capturing the variability in the data. When fitting EK-TOA, we suggest looking at these two types of residuals:

• **Standardized pre-fit difference residuals:** This shows how closely your CTCRW predictions line up with the observed data. This is simply the time-difference-of-arrival residuals after they have been standardized by the estimated residual covariance. Try to avoid movement parameters which result in overdispersion here. In our simulated study, overdispersion was associated with poor estimates of positioning error.
• **Smoothed residuals:** Here, the residuals are measured after smoothing has been applied.

```
plot(residuals(ektoa, type = 1), breaks = 200, xlim = c(-6, 6))
```

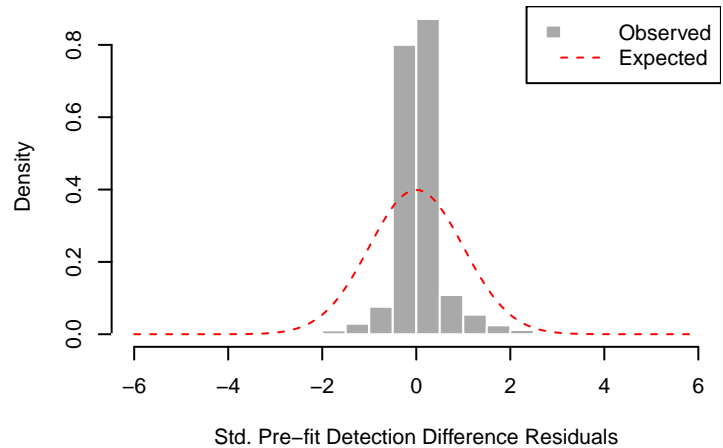

```
plot(residuals(ektoa, type = 2), breaks = 200, xlim = c(-0.01, 0.01))
```

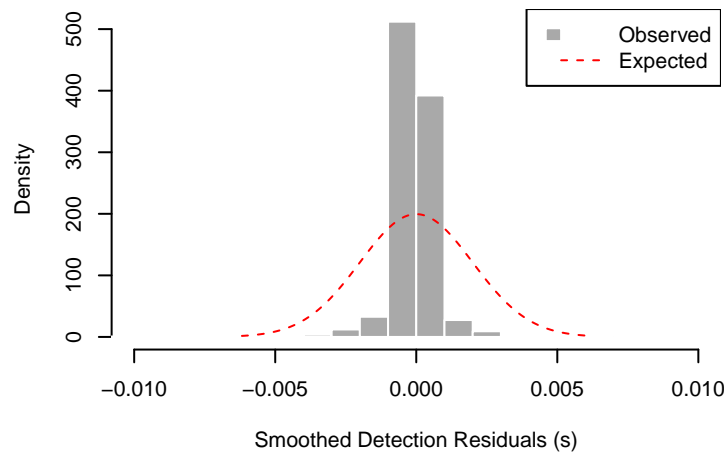

Examining these residual distributions can help inform your interpretation of the 95% confidence ellipses for the position estimates. Note that our residual distributions are underdispersed here. This means our confidence intervals are conservative, overestimating the true positioning error.

`state_covariance()` will return an array holding the covariance matrices for each state. These covariance matrices are used to create a confidence polygon — a union of the confidence ellipses for each state position estimate. `confidence_ellipse()` is a convenience function for creating these polygons.

```
ply_conf <- confidence_ellipse(
  ektoa, p = 0.95, circle_points = 20, dTolerance = 0.1) ► st_sfc(crs = crs)
```

Setting a low number of `circle_points` and a high `dTolerance` can speed this up at the cost of a less precise polygon.

```
# --- Initialize plot
plot(pts_rec, pch = NA, xlab = "Easting (m)", ylab = "Northing (m)", axes = T)
# --- Draw 95% confidence polygon
plot(ply_conf, col = 'lightblue', add = T, border = F)
# --- Draw study area
plot(ply_site, add = T)
# --- Draw receivers
plot(pts_rec, add = T, pch = 17, col = 'red')
# --- Draw fitted track
```

```
lines(state_position(ektoa), col = 'blue')
```

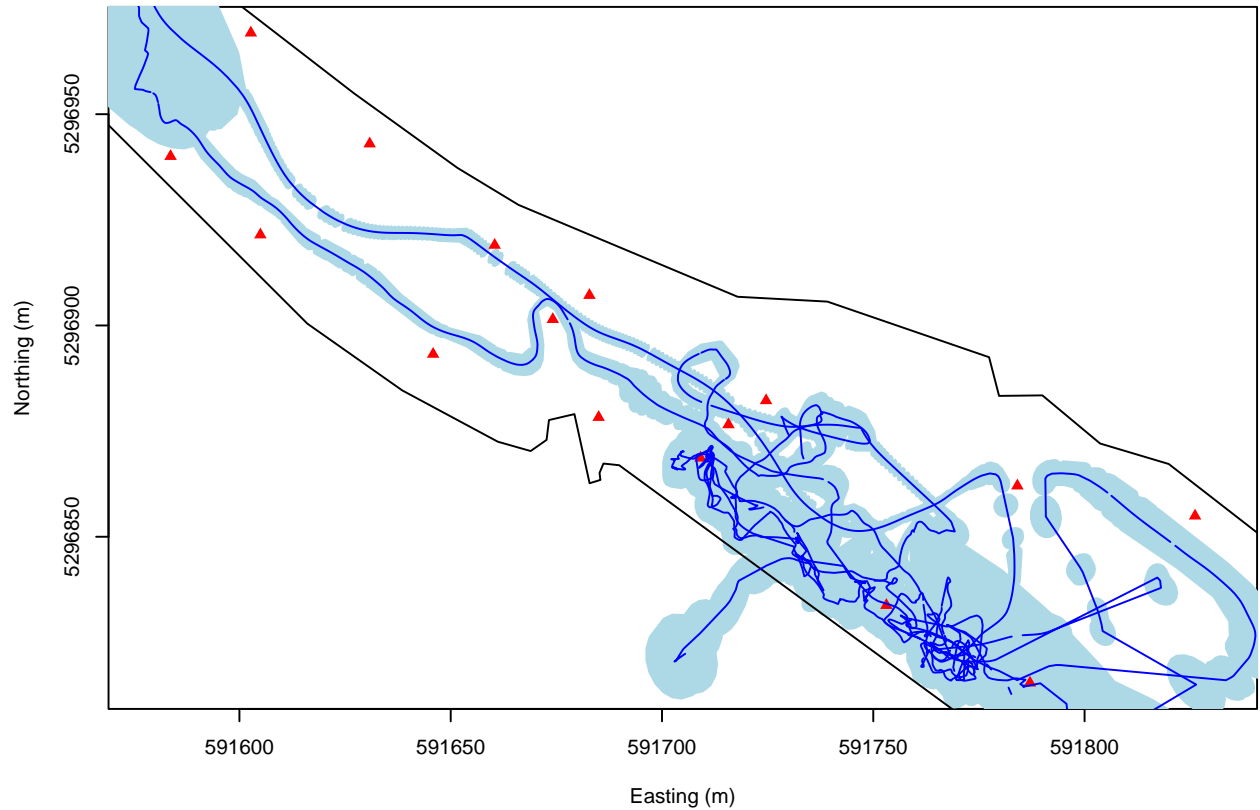

Finally, positioning error can be visualized as a histogram of some accuracy measure. Here, positioning accuracy can be defined as the semi-major axis of the 95% confidence ellipse drawn around a position estimate. That is, the length of the line drawn from the center of each confidence ellipse to its farthest edge.

```
pos_accr <- position_accuracy(ektoa = ektoa, p = 0.95)
pos_accr_hst <- hist(pos_accr[pos_accr < 50], plot = F,
  breaks = 50)
plot(pos_accr_hst, col = "darkgrey", border = "white",
  main = NULL,
  ylab = "Density", xlab = "Position Accuracy (m)")
```

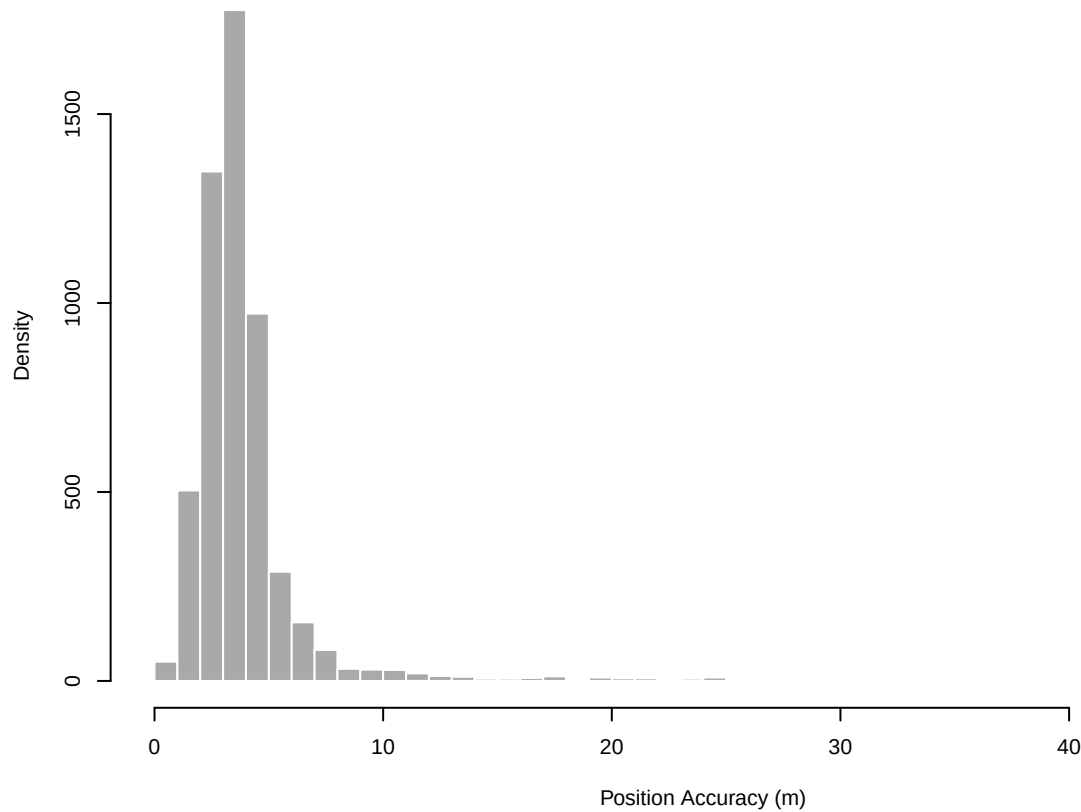

Barring some large outliers, the lion's share of the positioning sensitivity falls below 10 m, with a mode at roughly 4 m. These values can be viewed as a rough upper limit of the expected positioning error for a given position estimate.

### Simulating movement tracks

The accuracy of TOA positioning is highly dependent of the geometry of the receiver array. It's important to note that the EK-TOA model employs a linear approximation to this positioning model. Taken in consideration with the challenges of disentangling movement behaviour from TOA measurement error, simulated datasets are a useful tool for ensuring the behaviors of interest can be measured within a given study area.

The functions used to generate the simulated datasets found in the manuscript is given in `"./code/simulate_main.R"`. `make_track()` will generate a single-state CTCRW movement track that stays within a defined study area.

```
# --- Load simulation functions
source("./code/simulate_main.R")
# --- Set simulation parameters
# Tag emission intervals
sim_delta <- rep.int(15, times = 299)# (s)
# CTCRW movement parameters
sim_phi <- 0.2# (m/s)
sim_t_05 <- 100# (s)
# --- Create Single simulated track
set.seed(500)# make random track reproducible
sim_states <- make_track(
  delta = sim_delta, phi = sim_phi, t_05 = sim_t_05, study_site = ply_clip)
sim_pos <- sim_states[,c("s.x", "s.y")]
```

```
# --- Plot resulting track
plot(ply_clip, axes = T)
lines(sim_pos, col = "blue")
```

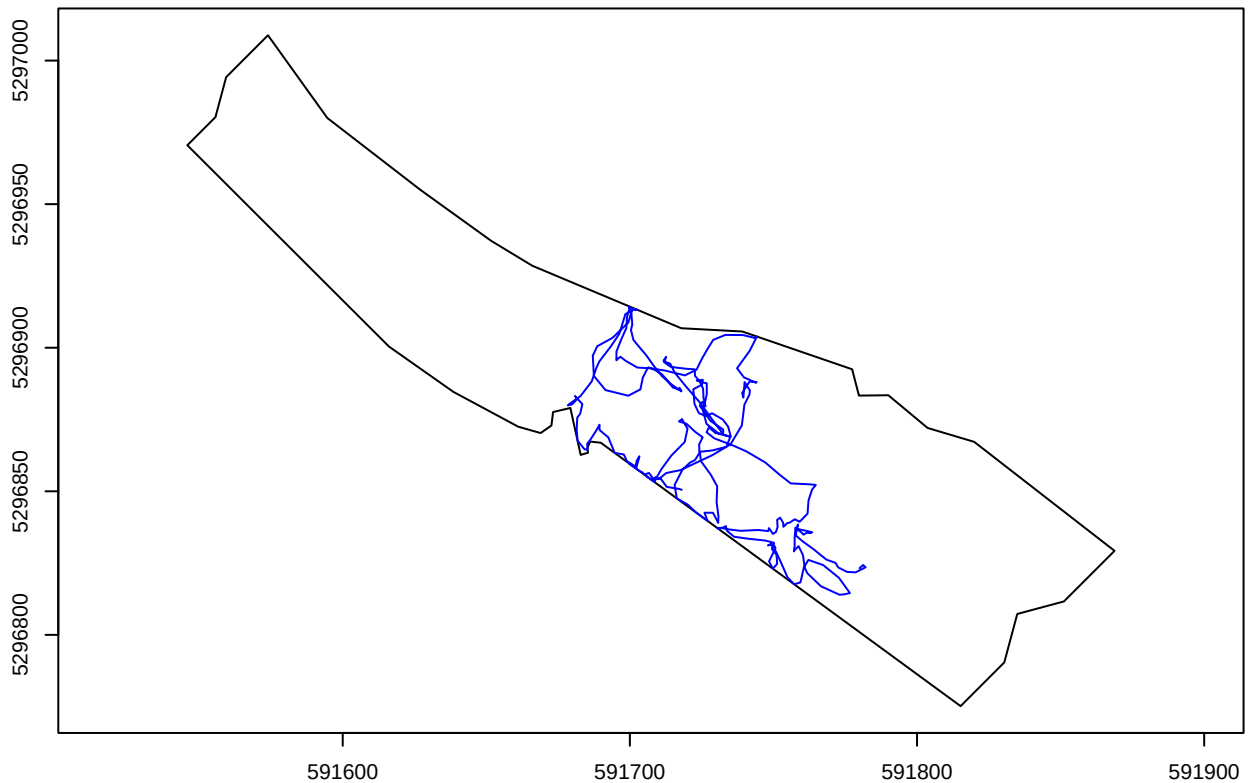

Next, simulate the time-of arrival values.

```
# ---Set TOA simulation parameters
# Set detection range
det_rng <- 150# (m)
# Set (within range) flat detection probability
det_p <- 0.7
# SD of Gaussian detection error
det_sigma <- 2e-3# (s)
# --- Simulate TOA Matrix
sim_toa <- make_toa(
  track = sim_pos, delta = sim_delta, # Track data
  pts_r = pts_rec, ply_site = ply_clip, # Study area
  det_rng = det_rng, det_p = det_p, det_sigma = det_sigma) # TOA parameters
# Remove emissions with no detections
sim_toa <- sim_toa[apply(sim_toa, 1, FUN = function(x) any(!is.na(x))),]
```

Finally, see how well the EK-TOA model fits to the simulated track.

```
# --- Fit EKTOA model to simulated TOA data
ektoa_2 <- eKalTOA(
  toa_matrix = sim_toa, r = mat_rec, c = 1500, d_max = 150,
  init_s = init_pos, init_s_sd = init_pos_sd,
  sigma_det = det_sigma,
  alpha = alpha, beta = beta,
  smooth = T)
```

```
# --- Plot EK-TOA fitted vs true tracks
# Study site
plot(ply_clip, col = "lightblue1", axes = T, border = F)
plot(ply_site, add = T)
plot(pts_rec, add = T, col = 'red', pch = 17)
# Simulated track
lines(sim_pos, col = 'blue', lt = 2)
# Fitted positions
lines(state_position(ektoa_2), col = 'black')
legend(x = "topright", legend = c("EK-TOA fit", "Sim.trk."),
      lty = c(1,2), col = c("black", "blue"))
```

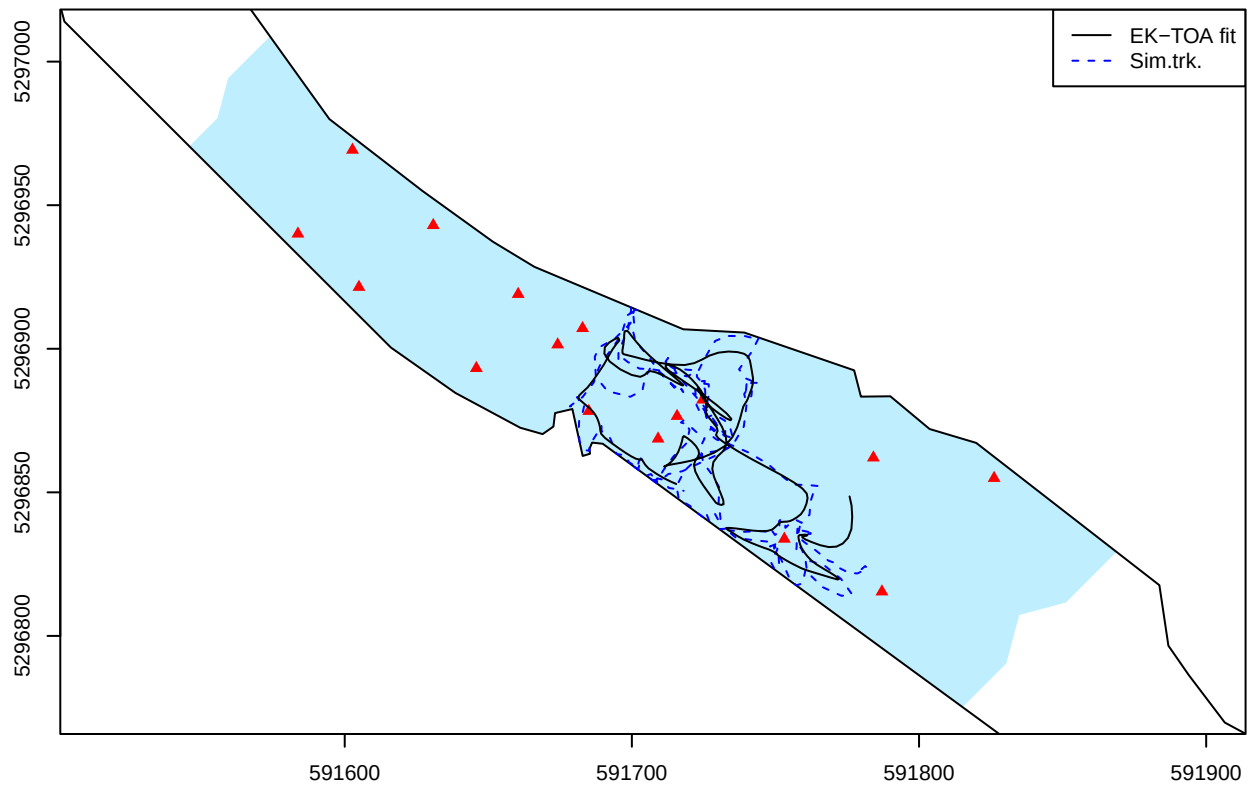
