## Supplementary material for "An extended Kalman filter for large-volume path positioning of aquatic animals within acoustic telemetry arrays": Detection Error

### Supplementary Material: Measured detection error

For the case study, the detection error of the array was measured using sentinel tags. Each receiver held a sentinel tag that logged the time of its emission events. For each combination of sentinel tag and receiver, the transmission latency was calculated — the recorded detection time at a receiver subtracted by the recorded emission time of the tag. Per each pair, these transmission latencies then had the expected transmission time (given a velocity of $1500 \text{ m s}^{-1}$) subtracted from them. The resulting values are the sentinel tag detection error: The observed minus expected detection times.

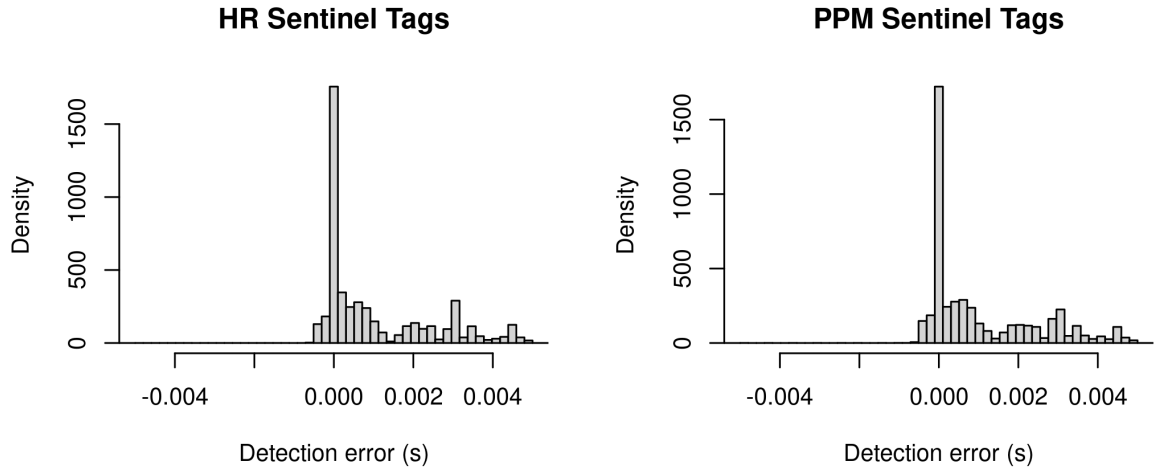

The above histograms show the pooled sentinel tag detection error across all tag/receiver combinations. Large positive outliers are likely the result of reflected transmissions. This error data was used as a guide to set the observation error parameter, $\sigma_{\text{det}}$, in the real and simulated case studies. These plots also served as a validation that the clock synchronization was applied correctly. Note that with a sound speed of $1500 \text{ m s}^{-1}$, an error of $1 \text{ ms}$ translates to a spatial positioning error of $1.5 \text{ m}$.
