## Supplementary material for "An extended Kalman filter for large-volume path positioning of aquatic animals within acoustic telemetry arrays": Pre-fit Residuals

Pre-fit residuals — also referred to as the innovation — are a useful diagnostic for recursive filters such as the Kalman filter. They are the difference between the model’s observed and expected measurements after the prediction step; in this case, the predicted time-of-arrival measures. When pre-fit residuals match poorly with their estimated covariance, this can be taken as an indication that more suitable filter parameters need to be chosen. A simple visual check can be done by plotting a histogram of the standardized pre-fit residuals — the residuals divided by their standard error — and verifying that the result falls along a standard Gaussian distribution.

In the context of EK-TOA, pre-fit residuals alone are not particularly informative. As telemetry tags are often set to emit on random intervals, the prediction step of EK-TOA has no information regarding when the next tag emission will occur. Hence, EK-TOA estimates of these absolute detection times from the prediction step are poor and the covariances of these estimates are rather large (owing to a large error for the initial emission time estimates, $\sigma_\epsilon$). The resulting standardized pre-fit residuals will typically result in a distribution far narrower than a standard Gaussian (Figure S1).

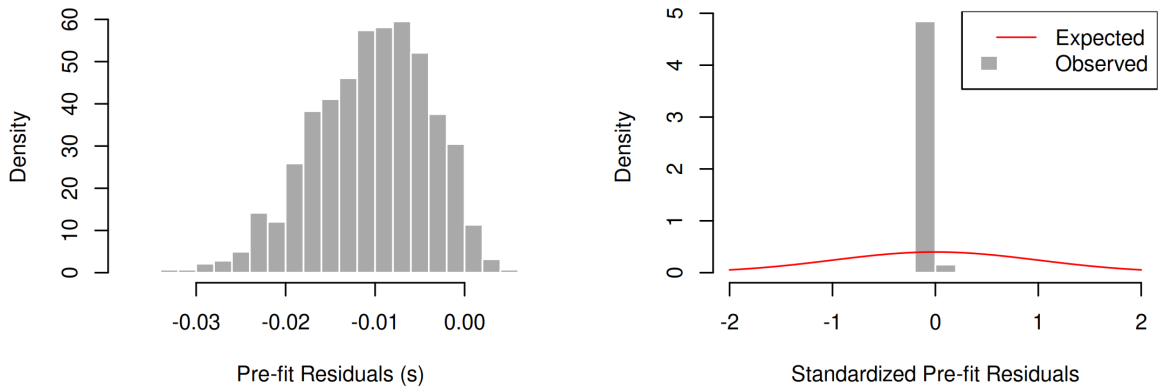

Fig. S1: Pre-fit residuals (time-of-arrival) returned from EK-TOA fitted to a single simulated track (track id: 10) with matching movement parameters. Despite using optimal movement parameters, the standardized pre-fit residuals are far smaller than their expected distribution.

These residuals, however, are highly correlated with each other across receivers for a given emission event. Pre-fit difference residuals can instead be taken by subtracting pre-fit residuals by the value returned from a selected receiver for each emission event. The residuals are now time-difference-of-arrival, rather than time-of-arrival values. The resulting standard error for these difference residuals will no longer be inflated by the large uncertainty of emission time estimates, providing a much more informative diagnostic (Figure S2).

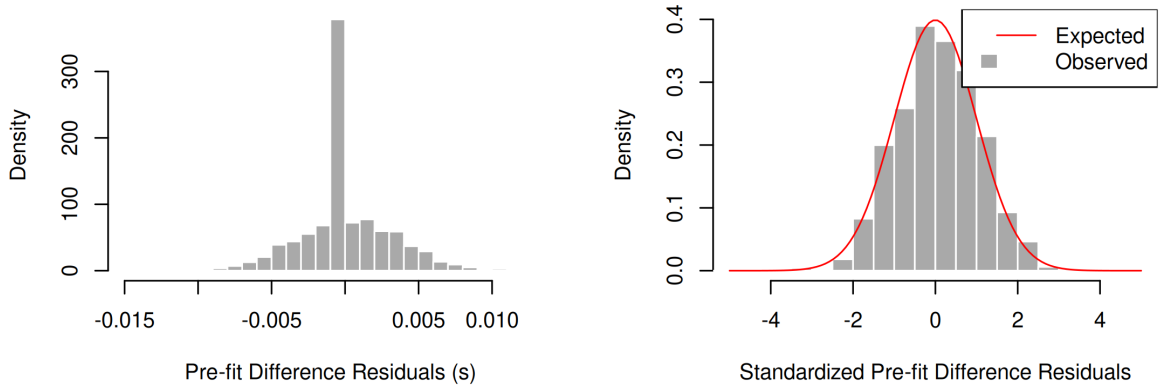

Fig. S2: Pre-fit difference residuals (time-difference-of-arrival) returned from EK-TOA fitted to a single simulated track (track id: 10) with matching movement parameters. Here, the uncertainty regarding the emission time has been removed from the residuals. As a result, the residuals match well with their expected distribution reflecting the optimal choice of movement parameters.

Figure S3 gives the pre-fit difference residuals for all EK-TOA models fitted in the simulated study. Those models using the same parameters as the simulated data (standard deviation of the stationary velocity, $\phi = 0.2 \text{ m s}^{-1}$; time until the velocity correlation falls below 0.05, $t_{05} = 100 \text{ s}$) gave values that matched well with a standard Gaussian. Generally, when $\phi$ or $t_{05}$ were set to low values the resulting distributions were overdispersed, indicating that the residuals were more extreme than EK-TOA expected.

Generally, it is advisable to choose movement parameters which cause the tails of the standardized residuals to lie within the expected standard Gaussian distribution. In the simulated study results, EK-TOA models which resulted in overdispersed residual distributions tended to also have poor estimates of both positioning uncertainty and absolute positioning accuracy. On the other hand, models fitted to large $\phi$ values gave more concentrated residuals with smaller tails. While these also resulted in lower position accuracy, the estimated positioning uncertainty remained well fitting. We suggest prioritizing trustworthy measures of positioning uncertainty over minimizing the absolute positioning accuracy.

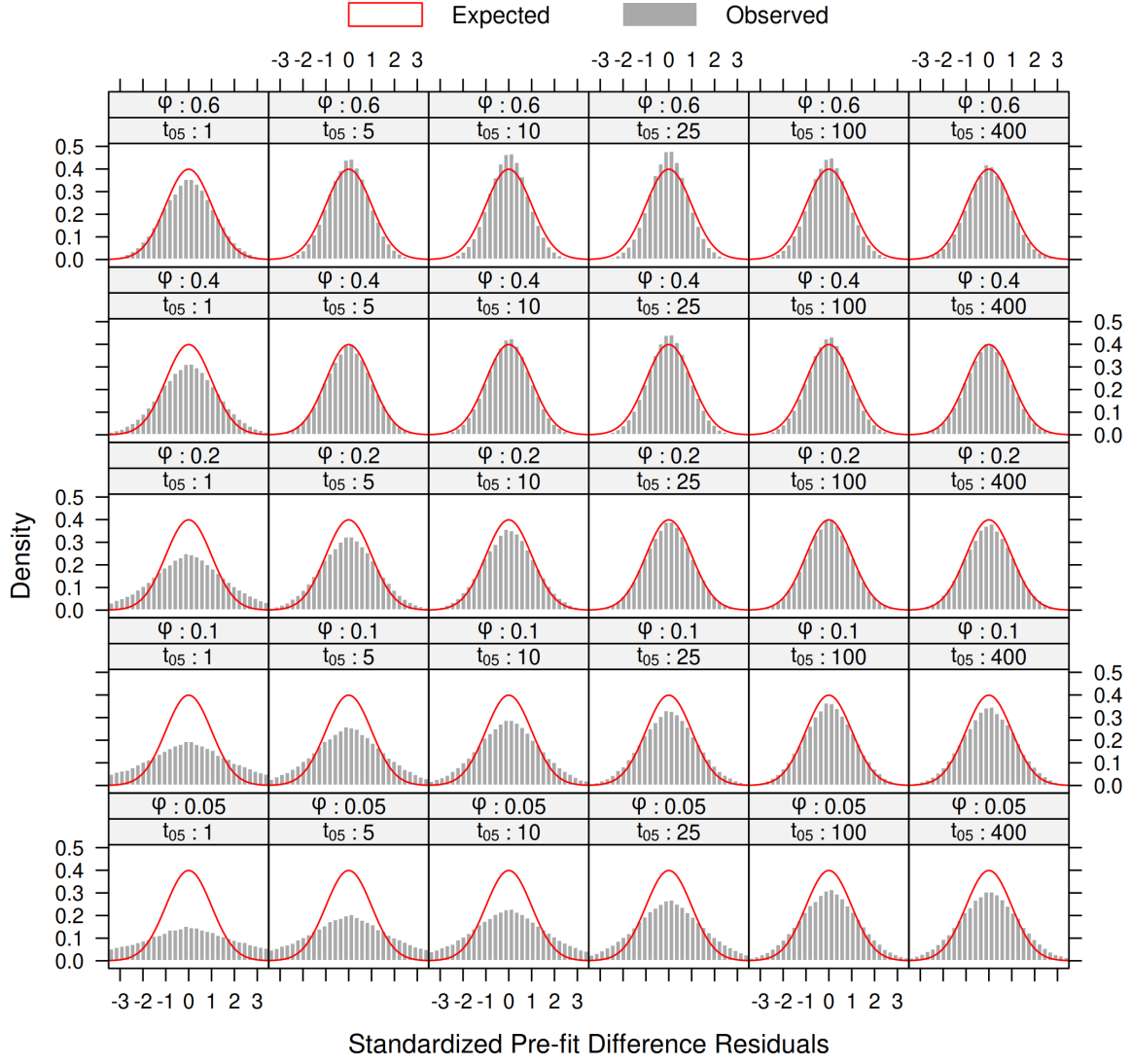

Fig. S3: Standardized pre-fit difference residuals returned from all EK-TOA models fitted across all simulated tracks. Gray bars are the observed residual distributions and the red lines give the expected standard Gaussian distribution. Deviations from the marked Gaussian lines can be used as an indicator of poor parameter selection. Each panel shows the pooled residuals across all EK-TOA fits for a given set of movement parameters. Generally, using too-small $\phi$ values resulted in overdispersed residual distributions that were correlated with poor estimates of positioning uncertainty.
