## Supplementary material for "An extended Kalman filter for large-volume path positioning of aquatic animals within acoustic telemetry arrays": Real case study figures

Figure S4 shows the estimated instantaneous velocity from EK-TOA after fitting to the real data case study. It is interesting to note the brief sections of track where the animals velocity was maintained at high values (at or above $0.84\text{ m s}^{-1}$) as it traversed large stretches of the study area. For the majority of the track, the tagged animal stayed at low velocities expressing relatively inactive behaviour states. These instantaneous velocity estimates may serve as a useful measure in identifying behaviour states or correlating the animals movements with the hydrodynamic environment of the study area (such as the flow velocity field).

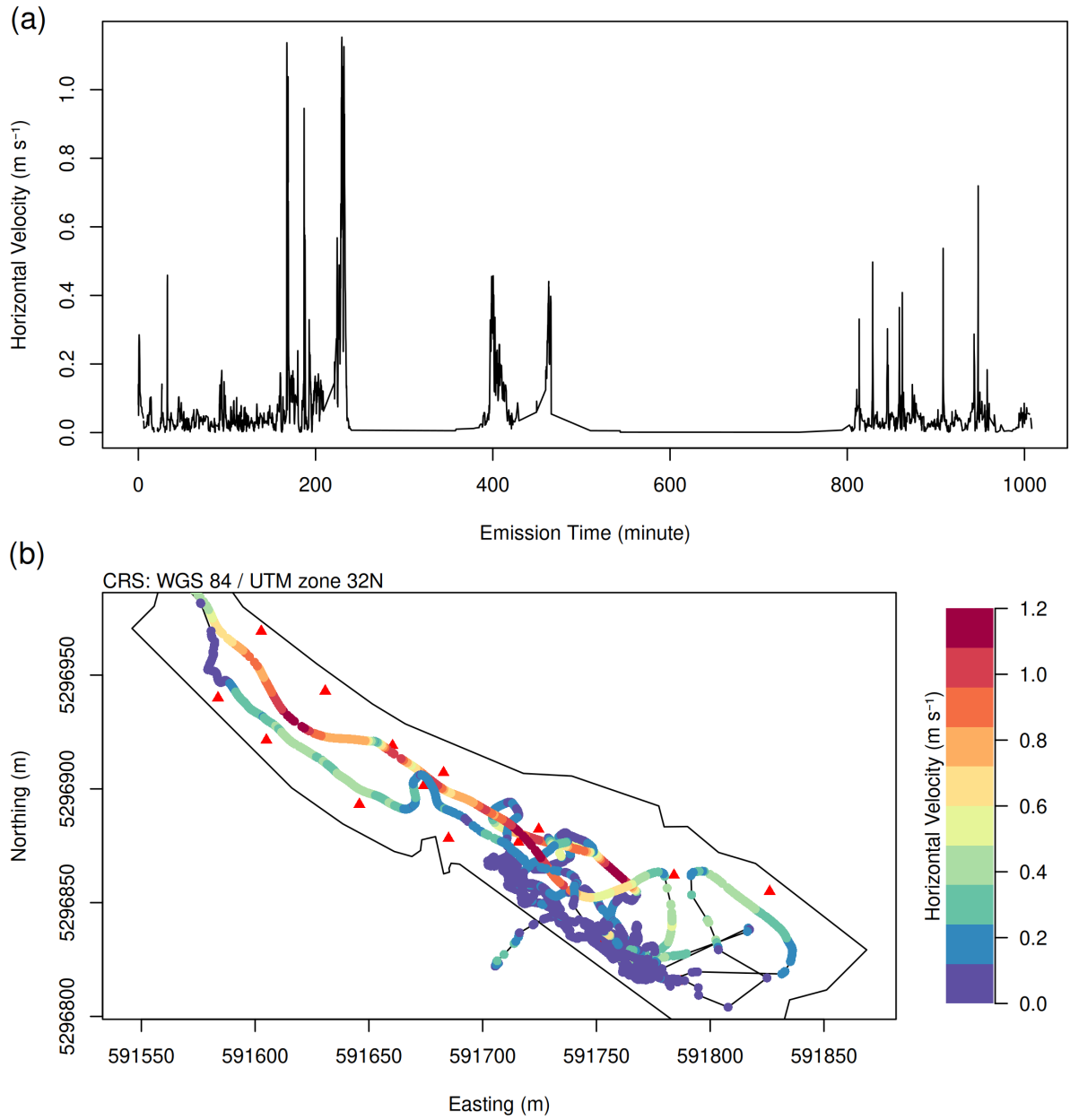

Fig. S4: Panel (a) shows the estimated instantaneous horizontal velocity over time for the real data case study. Panel (b) shows the fitted track overlaid with points colored according to the estimated instantaneous horizontal velocity.
