## Supplementary figures and images for "An extended Kalman filter for large-volume path positioning of aquatic animals within acoustic telemetry arrays"

### satellite.tif

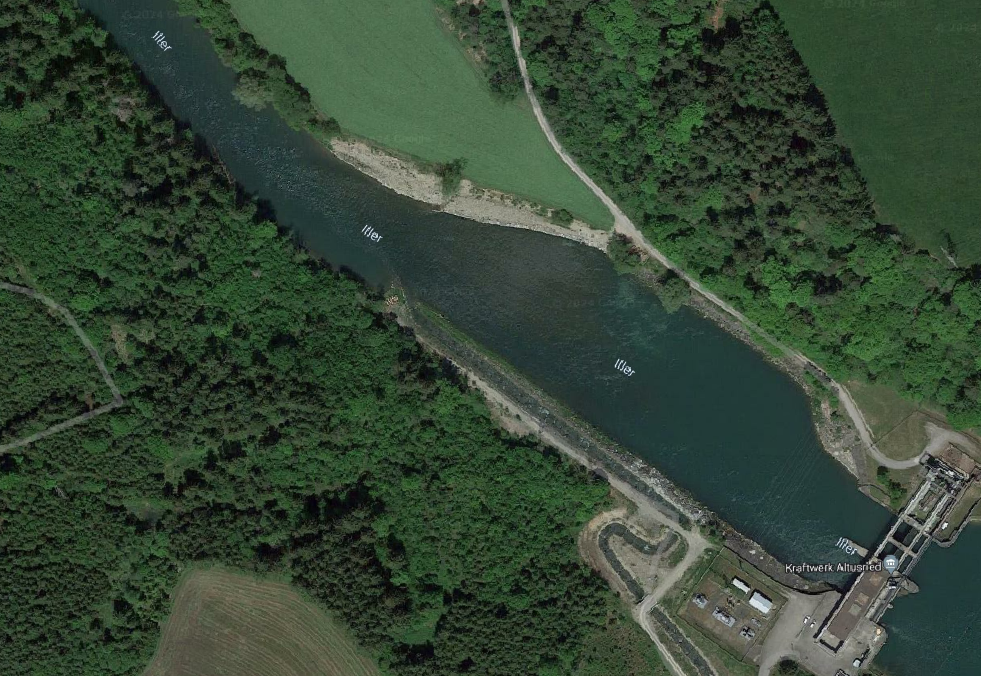
